# StructAgent: Orchestrating Cryo-EM Model Building and Refinement with a Multi-Agent LLM System

**DOI:** 10.64898/2026.05.18.725842

**Authors:** Xiaohu Guo

## Abstract

Building and refining cryo-EM atomic models often requires long, project-specific workflows that combine map inspection, prior structural knowledge, restraints, refinement, validation and expert review. Existing programs perform many individual operations, but coordinating them across iterative model-building sessions remains manual and difficult to audit. We present StructAgent, a user-guided multi-agent resource for cryo-EM model building and refinement. StructAgent couples a domain agent for literature-grounded structural reasoning with an execution agent that runs local software, tracks state, recovers from failures and records provenance. Expert approval gates control major model-changing actions. In three case studies, StructAgent refitted a 64-chain proteasome from an earlier template, audited 530 ribosomal metal-ion sites and guided a chemically ambiguous ligand fit in a folate-metabolism enzyme from ongoing work. These demonstrations show that agentic orchestration can convert modeling intent into auditable, reviewable software workflows while preserving expert control and final scientific judgment.

---

Cryo-electron microscopy (cryo-EM) has become central to the structural analysis of large, flexible and heterogeneous macromolecular assemblies. In many current projects, atomic model building no longer starts from a single map and an empty canvas. Structural biologists work with global and focused maps, predicted and homologous starting models, ligand or ion hypotheses, biochemical constraints and prior structural literature. Compared with typical crystallographic maps at comparable interpretive stages, cryo-EM maps are often noisier, have lower or more variable local resolution, and provide less direct evidence for side chains, ligands and ions.

Turning these inputs into a reliable model therefore requires adaptive sequences of fitting, rebuilding, restraint generation, refinement, validation and expert review. Mature programs support many individual steps, including automated model building [1–6], interactive flexible fitting [7] and reciprocal- or real-space refinement [5,8,9].

The remaining bottleneck is workflow coordination. Cryo-EM model building is rarely a linear run of one program: the user must choose tools, set parameters, inspect intermediate outputs, recover from failures, compare alternative strategies and document why each step was taken.

This burden is especially large for multi-chain assemblies, flexible conformational states, focused refinements, difficult ligand chemistry and large-scale local validation tasks. The unmet need is therefore not only to generate coordinates, but to make complex model-building workflows easier to explore, audit and repeat. A structural biologist may want to test whether domain-wise fitting improves a flexible assembly, whether a different refinement route better preserves geometry, whether a ligand orientation is chemically defensible, or whether hundreds of metal-ion sites should be re-evaluated under explicit coordination assumptions.

Large language models (LLMs) provide a possible interface for this type of workflow orchestration because they can translate natural-language intent into plans, commands and tool calls [10–18]. Related agent systems have begun to coordinate scientific tools in chemistry, drug discovery and protein-science settings [19–23]. General-purpose LLM use, however, is not sufficient for structural biology. Atomic model building is a high-consequence scientific task: hallucinated commands, unsupported interpretations, undocumented parameters, lost context or unreviewed model changes can compromise reproducibility. Long cryo-EM sessions add a further challenge because tool logs, file paths, errors and validation outputs can accumulate over hours or days. An LLM-based system for cryo-EM therefore needs a constrained operating model, not only conversational fluency.

We developed StructAgent as a user-guided orchestration resource for cryo-EM model building and refinement. StructAgent separates scientific reasoning from execution management through two cooperating agents. Maria, the domain agent, evaluates strategies using a curated structural-biology knowledge base, literature retrieval and project memory. Annika, the execution agent, translates approved strategies into local software actions, manages files and state, monitors outputs, diagnoses failures and records provenance. Both agents operate through editable skill documents that encode tool use, workflow logic, validation checks and recovery procedures for programs such as ChimeraX, ISOLDE, PHENIX, Coot, Servalcat/REFMAC5 and gemmi. The trust model is deliberately conventional: established software performs the calculations; skills and literature constrain reasoning; major model-changing actions require expert approval; and runs leave logs, metrics, inputs, outputs and rationale for later review.

The operational unit of the system is the skill. Skills are readable protocol documents rather than hidden prompts: they specify expected inputs, command patterns, validation metrics, common failure modes and recovery actions. This makes the resource inspectable and editable by the same researchers who rely on it. The system can therefore accumulate workflow knowledge without turning it into an opaque automation layer; a user can review how a tool was called, why a recovery action was chosen and which parts of the protocol should be revised after a failed or successful run.

We evaluated StructAgent using three representative cryo-EM task classes. In a 64-chain proteasome example, StructAgent coordinated domain-aware refitting, restraint generation, flexible fitting, refinement and validation starting from an earlier structural template. In a 1.55 Å ribosome example, it built and executed a systematic audit of 530 metal-ion sites and refined 75 user-approved corrections with tailored restraints. In a ligand-fitting example, it combined local density fitting with literature-grounded reasoning to choose the orientation of a folate-derived ligand in an ambiguous binding site. These examples are case studies, not formal benchmarks or universal protocols. They test whether an agentic system can coordinate multi-tool cryo-EM workflows, recover from execution failures and provide evidence packages for expert review. StructAgent is not a coordinate-prediction algorithm or an autonomous substitute for structural biology expertise; it is an auditable layer that helps experts convert modeling intent into reviewable computational workflows.

## Results

### System overview

StructAgent is a conversational workflow system for cryo-EM model building and refinement (Fig. 1). The user communicates in natural language through standard messaging applications, while all structural-biology software runs locally. Maria provides literature-grounded domain reasoning and validation criteria. Annika manages local execution, retry logic, workflow state, validation output and provenance capture. The two-agent design is intended to keep execution-heavy context separate from scientific reasoning during long sessions; inter-agent consultations are logged through the A2A channel and described in Online Methods. StructAgent uses editable skill protocols (Supplementary Table 1), a curated knowledge base (Supplementary Table 2), established structural-biology software, approval gates and session logs as its audit framework. The output is not an autonomous final structure but a reviewable package of models, maps, metrics, logs and rationale from which the structural biologist chooses the next action.

**Fig. 1.**
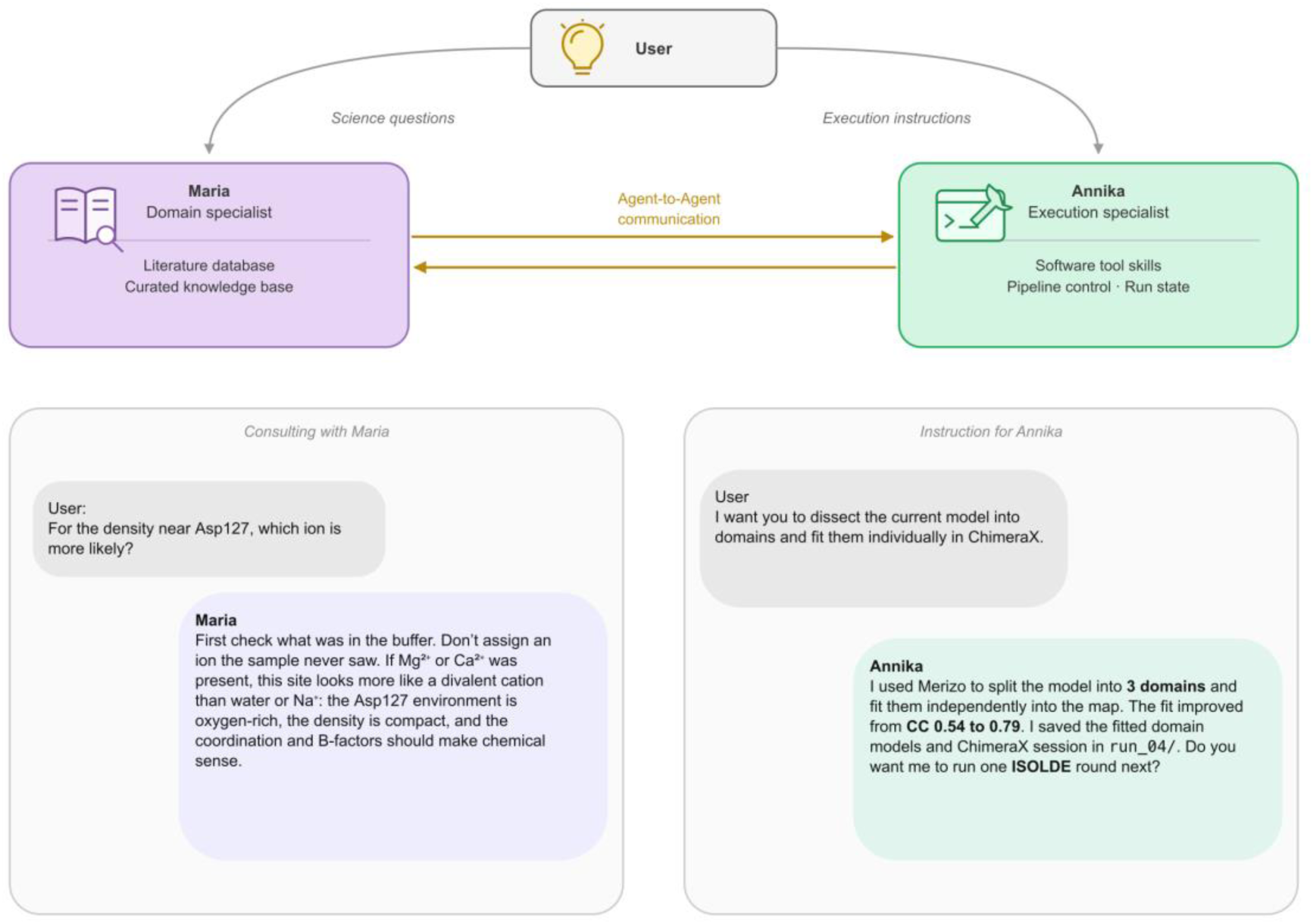
StructAgent separates structural-biology reasoning from local software execution. The user supplies scientific intent, constraints, approvals and final interpretation. Maria provides literature-grounded reasoning and validation criteria; Annika manages local execution, retries, logs and provenance. The audit framework combines curated literature and skills, established structural-biology software, approval gates, validation output and provenance records. The intended output is a reviewable evidence package for expert decision-making.

### Representative cryo-EM task classes

The three examples below stress different aspects of StructAgent: large-assembly rebuilding, systematic local validation and chemically informed ligand fitting. They demonstrate rapid, auditable exploration of model-building ideas rather than benchmark superiority over expert manual workflows.

### Example 1: whole-assembly rebuilding of the 26S proteasome

We first tested whether StructAgent could coordinate a whole-assembly rebuild rather than a single local fitting decision. The target was the human 26S proteasome map EMD-8332 at 3.8 Å, with 64 chains, 32 unique subunits and 19,957 residues in the rebuilt model. As the starting scaffold, we used PDB 5GJR, an earlier related 26S proteasome model that provided chain topology but was a poor rigid-body fit to this map. Deposited PDB 5T0C, the expert-refined model for EMD-8332, was held out from the workflow and used only as a post hoc reference.

Fig. 2a summarizes the division of work. The user supplied the input files and rebuilding goal; Maria analyzed the inputs, wrote the staged plan and later reviewed the refinement results; Annika executed the approved workflow. The task required domain definition, chain anchoring, restraint generation, flexible fitting, reciprocal-space refinement, real-space refinement and validation across all 64 chains. Annika generated the chain-specific fitting scripts using Merizo [24] for domain segmentation and ProSMART [25] restraints derived from AlphaFold [26] models, then ran the workflow with ChimeraX, ISOLDE, Servalcat/REFMAC5, PHENIX, gemmi and supporting scripts.

**Fig. 2.**
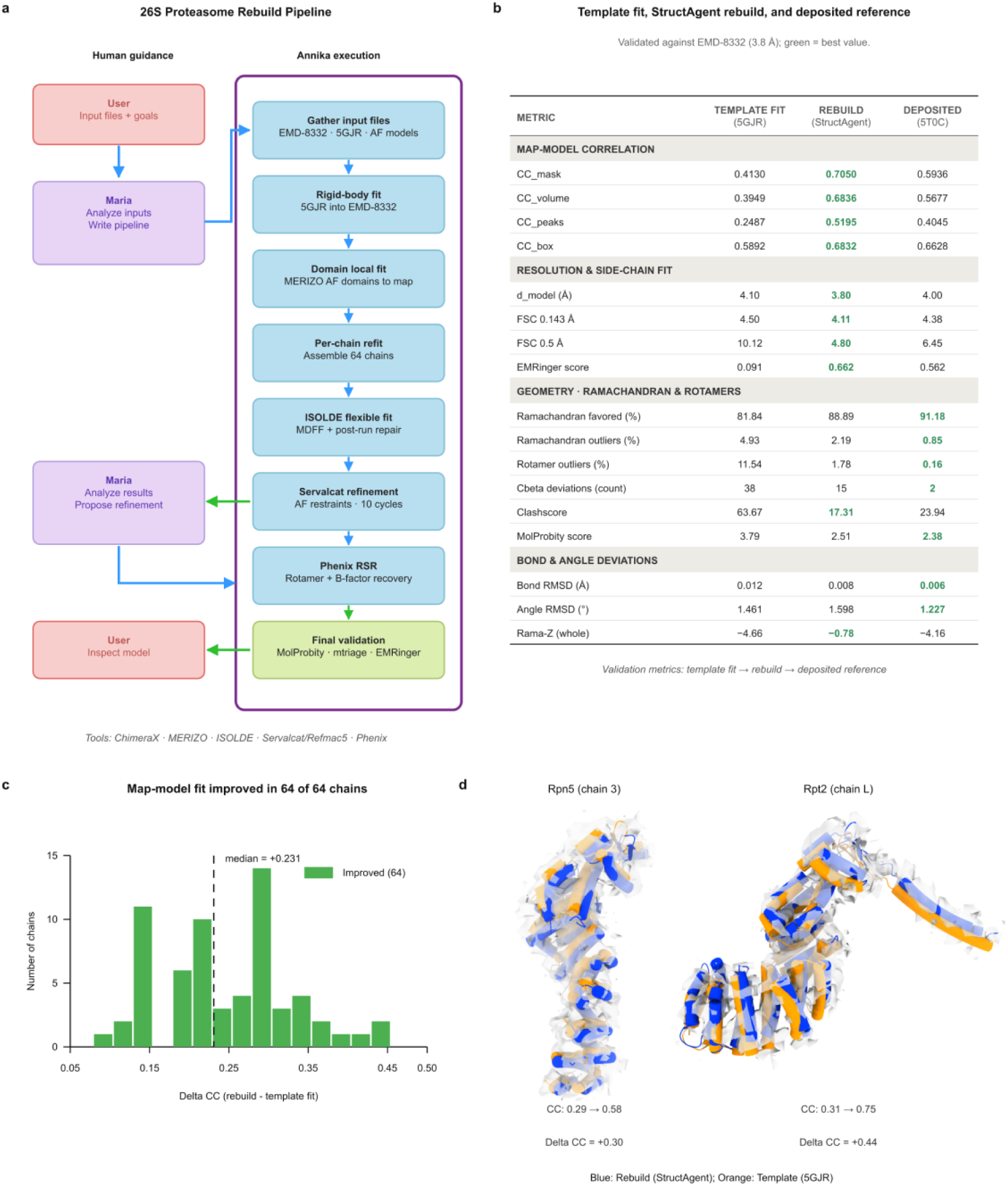
Whole-assembly rebuilding of the 26S proteasome from an earlier structural template. **(a)** StructAgent pipeline for refitting PDB 5GJR into EMD-8332. Recovery events are annotated. **(b)** Validation comparison among the rigid-body 5GJR template fit, the StructAgent rebuild and deposited 5T0C, all evaluated against EMD-8332 (3.8 Å full map; half-maps not used). Metric direction: higher is better for CC, EMRinger and favored percentages; lower is better for outliers, clashscore, RMSD and FSC resolution; Rama-Z is interpreted relative to zero. **(c)** Per-chain map-correlation improvement for all 64 chains relative to the template fit (mean ⊗CC = +0.24, median = +0.23). **(d)** Local density comparisons for Rpn5 (chain 3, ⊗CC = +0.30) and Rpt2 (chain L, ⊗CC = +0.44). Orange, 5GJR template fit; blue, StructAgent rebuild.

The workflow did not succeed by running a fixed script end to end. An initial AlphaFold-only assembly route produced unstable inter-domain geometry, so Maria re-planned the rebuild around 5GJR as a topology scaffold and AlphaFold monomers as domain and restraint references. Annika then rigid-fit 5GJR into the map, refit each chain through its best-matching domain anchor, assembled the full complex and carried the model through ISOLDE flexible fitting, Servalcat/REFMAC5 refinement and PHENIX real-space refinement. During this process Annika recovered five tracked execution failures, including force-field parameterization gaps, a hydrogenation-induced coordinate-frame shift, post-simulation chain drift, a case-insensitive chain-file collision on macOS and B-factor inflation after refinement (Supplementary Table 3).

The reported StructAgent rebuild converted a weak template fit into a strong expert-handover model (Fig. 2b). The figure compares three states: the rigid-body 5GJR template fit, the StructAgent rebuild and the deposited 5T0C reference, all validated against EMD-8332.

CC_mask increased from 0.413 to 0.705, EMRinger from 0.091 to 0.662, rotamer outliers decreased from 11.54% to 1.78% and clashscore from 63.67 to 17.31. Under the validation pipeline used here, the rebuild also exceeded deposited 5T0C on density-agreement metrics, including CC_mask (0.705 versus 0.594), EMRinger (0.662 versus 0.562) and FSC = 0.5 (4.80 versus 6.45 Å). This is workflow evidence, not a claim that the rebuilt model supersedes 5T0C: validation used the deposited full map because half-maps were unavailable, and stronger density-fit scores may reflect overfitting risk or differences in modern restraints.

The comparison also defined the boundary of the automation. Local geometry improved substantially from the 5GJR template, but deposited 5T0C remained better on rotamer outliers (0.16% versus 1.78%), Ramachandran outliers (0.85% versus 2.19%), C® deviations and bond/angle RMSD. The rebuild’s better Rama-Z whole score (−0.78 versus −4.16 for 5T0C) suggests that some validation differences reflect newer refinement libraries, but it does not remove the need for expert inspection of local side-chain choices at 3.8 Å.

The per-chain analysis in Fig. 2c shows that the improvement was broad rather than driven by one selected region. All 64 chains improved in map agreement relative to the rigid-body 5GJR fit, with mean ⊗CC = +0.24 and median ⊗CC = +0.23. The local examples in Fig. 2d make the same point visually: Rpn5 improved from CC 0.29 to 0.58, and Rpt2 from 0.31 to 0.75. In the post hoc comparison with deposited 5T0C, the rebuild had higher per-chain CC for 61 of 64 chains, with only small regressions in the remaining three.

The endpoint was therefore expert handover, not autonomous deposition. StructAgent had completed the coordination-intensive part of the rebuild, captured the failures and recovery steps, and produced a reviewable model package. The remaining work was the part that should remain with a structural biologist: targeted side-chain correction, local geometry review and final scientific judgment.

### Example 2: systematic metal-ion audit of a 1.55 Å ribosome

Metal-ion identities in cryo-EM models are often difficult to assign because Coulomb potential maps do not provide the anomalous signal available in X-ray crystallography. An experienced crystallographer can inspect a few suspected sites by comparing coordination distances, ligand types and expected geometry with tabulated values [27], but this process does not scale cleanly to large assemblies. PDB 8B0X, a 1.55 Å *E. coli* 70S ribosome [28], contains 530 deposited metal sites across 104 chains. Majorek et al. [29] had already reported four misassigned ions in this structure using the CheckMyMetal (CMM) validation server. We therefore asked whether StructAgent could turn classical coordination chemistry into a systematic, auditable audit, correction and restraint-generation workflow for every site, rather than a manual review of selected examples.

The first constraint came from the user rather than from the coordinates alone: Mg²⁺, K⁺ and Zn²⁺ were specified as candidate identities based on the buffer composition and known ribosomal chemistry. This experimental context limited the search to chemically plausible ions before any scoring was done. Maria then translated bond-valence-sum (BVS) parameter sets [30,31] together with Harding’s [27] metal-ligand coordination geometry into an executable specification, including R₀ parameters, expected coordination numbers, distance cutoffs, ligand-type checks and confidence grades. Annika implemented the audit with gemmi by extracting each coordination shell, scoring every site against the candidate ions and assigning grades from A to F. The full 530-site audit ran in a single session and separated plausible assignments from sites requiring review (Fig. 3a). After user review, 75 corrections were approved: 56 Mg²⁺ to K⁺, 10 Mg²⁺ to Zn²⁺, 2 Mg²⁺ to water and 7 K⁺ to water.

**Fig. 3.**
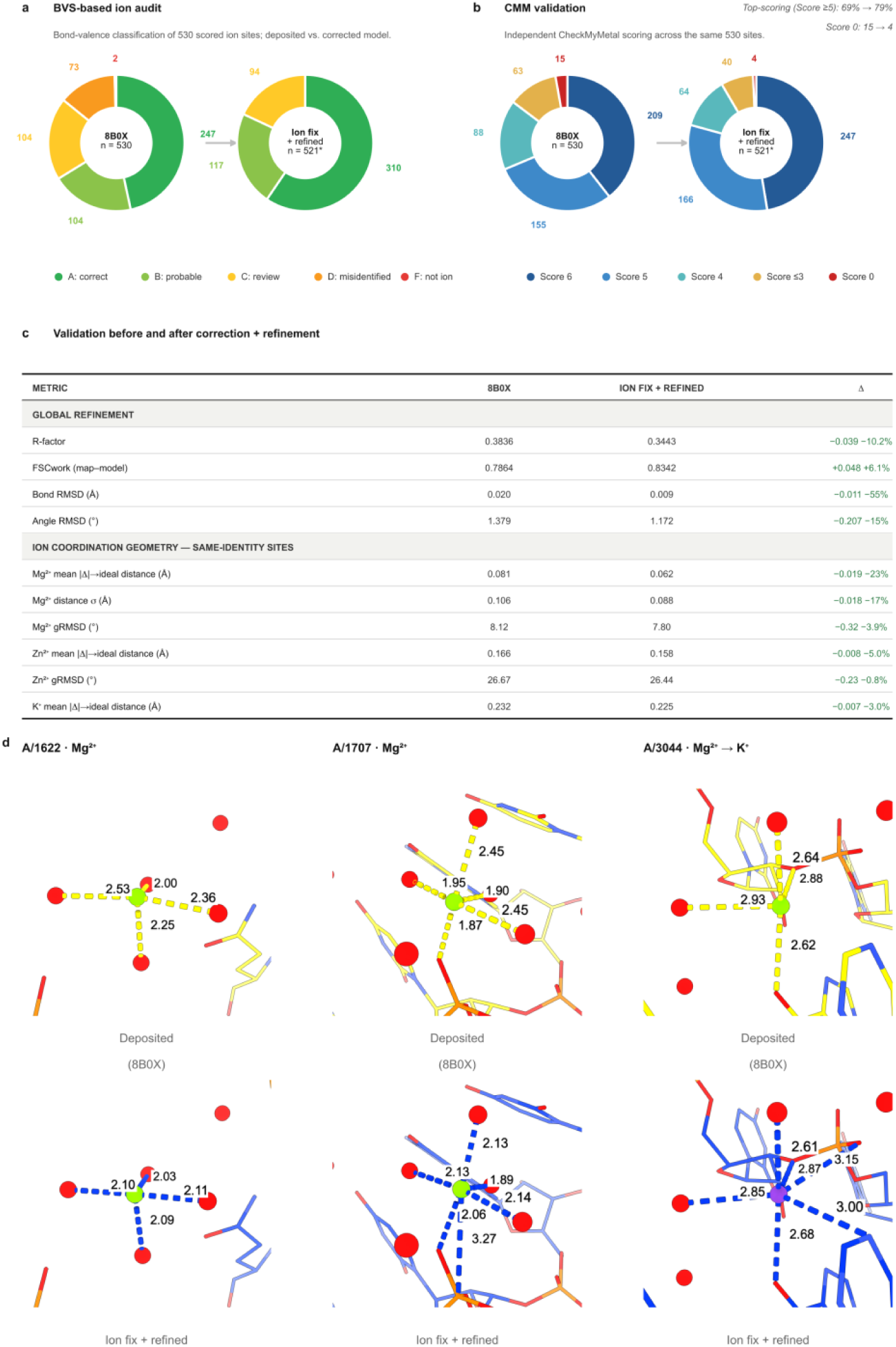
Systematic metal-ion audit and correction of a 1.55 Å cryo-EM ribosome. **(a)** BVS classification of all deposited ion sites in PDB 8B0X before correction (n = 530) and after correction and refinement (n = 521 ion sites; 9 original positions reclassified as water are excluded from corrected ion-score totals). Confidence grades A-F are assigned from BVS, coordination number and geometry. **(b)** Independent rescoring with the CheckMyMetal (CMM) validation server [32] across the original 530 positions; corrected-model ion-score totals exclude the 9 water-reclassified positions. Top-scoring sites increase from 69% to 79%, and score-0 outliers decrease from 15 to 4. **(c)** Global refinement and ion-coordination statistics before and after correction plus REFMAC5 refinement with bond-distance and angle-derived 1-3 distance restraints. **(d)** Three representative sites before (deposited 8B0X, top) and after (corrected + refined, bottom): A/1622 and A/1707 (correctly assigned Mg²⁺, with improved coordination geometry after refinement) and A/3044 (Mg²⁺ → K⁺ reclassification, one of four sites previously documented as misassigned by Majorek et al. [29]). BVS, bond-valence sum; CMM, CheckMyMetal; FSC_work, work Fourier shell correlation; gRMSD, geometric root-mean-square deviation; ⊗, corrected minus deposited.

The audit did not stop at a list of suspicious ions. Annika generated ion-specific coordination restraints for the corrected model and refined it with REFMAC5 through Servalcat. This step moved the workflow beyond what a validation server alone provides: the same literature-derived assumptions used to flag sites were converted into restraint files for refinement across more than 500 local environments. The most difficult recovery involved external angle restraints.

Servalcat’s chain-renaming scheme produced nonpolymer identifiers that REFMAC5 could not parse through the EXTE ANGL route. After five failed formulations, Maria recast the angle restraints as equivalent 1-3 distance restraints using the law of cosines, and Annika executed the refinement through EXTE DIST. This restraint cascade accounts for five of the eight Example 2 recovery events in Supplementary Table 3.

Independent CMM [32] rescoring supported the corrected model while keeping the comparison anchored to an established validation tool (Fig. 3b,c). CMM was applied to the original 530 deposited positions; after correction, the 9 positions reclassified as water were excluded from corrected ion-score totals, leaving 521 ion sites. Top-scoring CMM sites increased from 69% to 79%, score-0 outliers decreased from 15 to 4, R-factor improved from 0.3836 to 0.3443 and FSC_work improved from 0.7864 to 0.8342. The local examples in Fig. 3d show both kinds of outcome: retained Mg²⁺ sites with improved coordination geometry and a Mg²⁺ to K⁺ reassignment at A/3044, one of the sites previously documented by Majorek et al. [29].

This case is not presented as a universal metal-curation protocol or as a critique of the original deposition. The Zn²⁺ reassignments remain the least independently supported class, consistent with atypical or incomplete peripheral coordination environments. The demonstration is instead that StructAgent could combine user-supplied experimental context, literature-grounded BVS rules, large-scale coordinate analysis, restraint generation, refinement and independent validation into one reviewable workflow. Conventional tools scored sites or refined coordinates; the agentic contribution was connecting those operations into a chemically reasoned strategy at a scale that is impractical to perform by hand.

### Example 3: ligand fitting with literature-grounded orientation reasoning

Ligand fitting in cryo-EM often requires more than maximizing map correlation. At typical local resolutions, ligand density can be incomplete, flexible groups may be weak or disordered, and multiple chemically different poses can give plausible local fits. We therefore tested whether StructAgent could use prior structural knowledge to guide ligand placement while still keeping the decision auditable. The target was a bifunctional folate-metabolism enzyme from an ongoing cryo-EM study. The unliganded 3.2 Å focused map lacked 5-formimidoyltetrahydrofolate (CHEBI:57456), an approximately 18 Å folate derivative, at two sites. One site, the transferase active site, had a direct 1.7 Å X-ray homolog (PDB 1QD1, ligand FON). The second site, at the inter-domain interface, had no homologous ligand pose and showed elongated density compatible with two orientations.

The executable workflow is summarized in Fig. 4a. Annika routed the task through existing tools for restraint generation, homolog transfer, flexible fitting, real-space refinement and validation, and recovered an ISOLDE/OpenMM template failure by generating a USER_LIG force-field file (Supplementary Table 3). The figure carries the tool sequence; the central result is the decision logic for the inter-domain interface site.

**Fig. 4.**
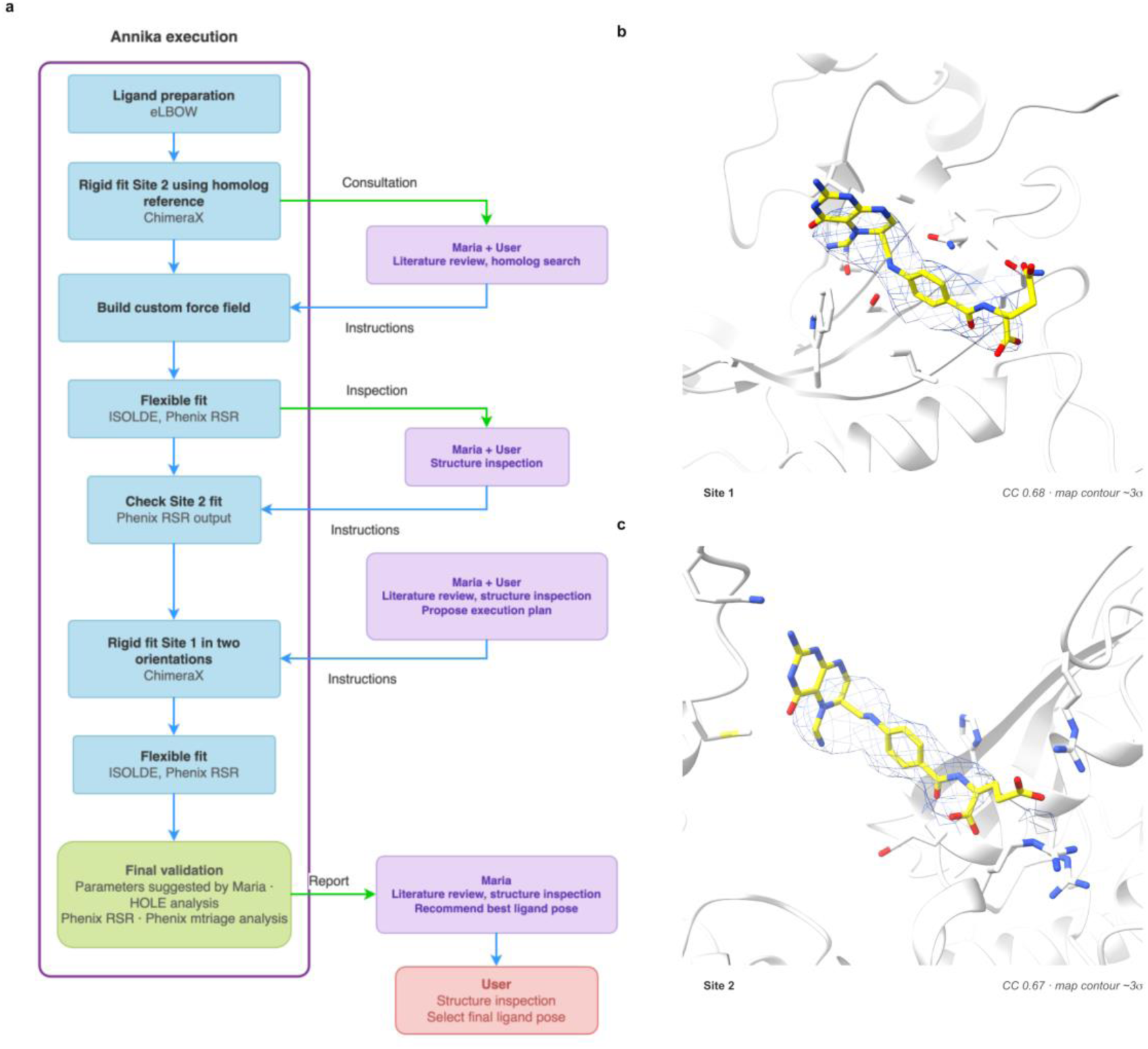
Folate-ligand fitting into cryo-EM density with scientific reasoning. (a) Workflow for fitting 5-formimidoyltetrahydrofolate into two sites in an ongoing-work enzyme. Annika’s execution steps are shown with Maria/user consultation points. (b) Site 1, the inter-domain site without a homologous ligand pose, after orientation selection, ISOLDE fitting and PHENIX refinement; per-ligand CC = 0.68. (c) Site 2, the transferase active site fitted by homolog transfer from PDB 1QD1/FON and refined with the two-site model; per-ligand CC = 0.67. The glutamate tail shows weak density consistent with conformational disorder at approximately 3.2 Å local resolution. The map and model are from an ongoing biological study and will be deposited separately.

Maria evaluated the ambiguous site by combining residue chemistry, folate-binding literature and substrate-channeling logic, rather than by asking which orientation gave the highest density score alone. She recommended the pterin-to-chain-C orientation because Tyr419 could stack the pterin ring, Arg424, Arg436 and Asp423 could support pterin recognition, and the Lys193-His197-Arg198-Arg381 positive cluster faced the glutamate carboxylates. ChimeraX fitmap supported this choice but did not by itself settle it: the selected orientation had z_mean 7.35 versus 6.06 for the alternative (⊗z_mean = 1.29). At Maria’s request, Annika then ran a per-group comparison on unrefined seed poses; the selected orientation was better in every ligand group, including pterin-ring CC 0.19 versus 0.04 and formimidoyl z-score 9.27 versus 4.68.

Maria also shaped the validation endpoint. The first combined PHENIX refinement placed both ligands, but diagnostics showed frozen ligand B-factors and pterin-ring planarity distortion. Maria advised a single combined PHENIX pass with individual isotropic ADP refinement and custom pterin planarity restraints, while explicitly rejecting manual B-factor rescaling and over-restraining of the flexible glutamate tail. This raised CC_mask from 0.7997 to 0.8194, improved the inter-domain interface ligand CC from 0.5848 to 0.6785 and gave the transferase active-site ligand CC of 0.6713 (Fig. 4b,c).

The final evidence package reported both support and uncertainty. HOLE identified a continuous channel between the FT and CD sites with a minimum radius of 6.09 Å, consistent with the selected orientation. The Site 1 glutamate tail remained weak, with local CC approximately 0.09, consistent with a surface-exposed flexible tail at 3.2 Å; this was reported rather than forced into density. This case therefore highlights the role of Maria: conventional software generated poses, scores and refined coordinates, but the decisive contribution was literature-grounded chemical interpretation, diagnostic review and restraint choice.

### Cross-case summary

Across the three case studies, StructAgent coordinated workflows using ChimeraX, ISOLDE, PHENIX/eLBOW, Servalcat/REFMAC5, gemmi, Coot and custom validation scripts; recovered all 14 tracked tool-level failures; and produced reviewable evidence packages containing models, maps, metrics, logs and rationale. The agent architectures supporting this division of domain reasoning and execution management are summarized in Fig. 5. Fig. 6 illustrates why this separation matters during long workflows, where execution history can otherwise compete with domain-level reasoning. The corrected failure distribution is 5/5 for the proteasome rebuild, 8/8 for the metal-ion audit and 1/1 for ligand fitting (Supplementary Table 3). Most recoveries were handled by Annika without user intervention; the hardest cases required Maria’s domain reasoning or explicit user approval for model-changing decisions. The case studies also illustrate how successful recovery patterns can be distilled into future skill updates rather than remaining as one-off troubleshooting episodes.

**Fig. 5.**
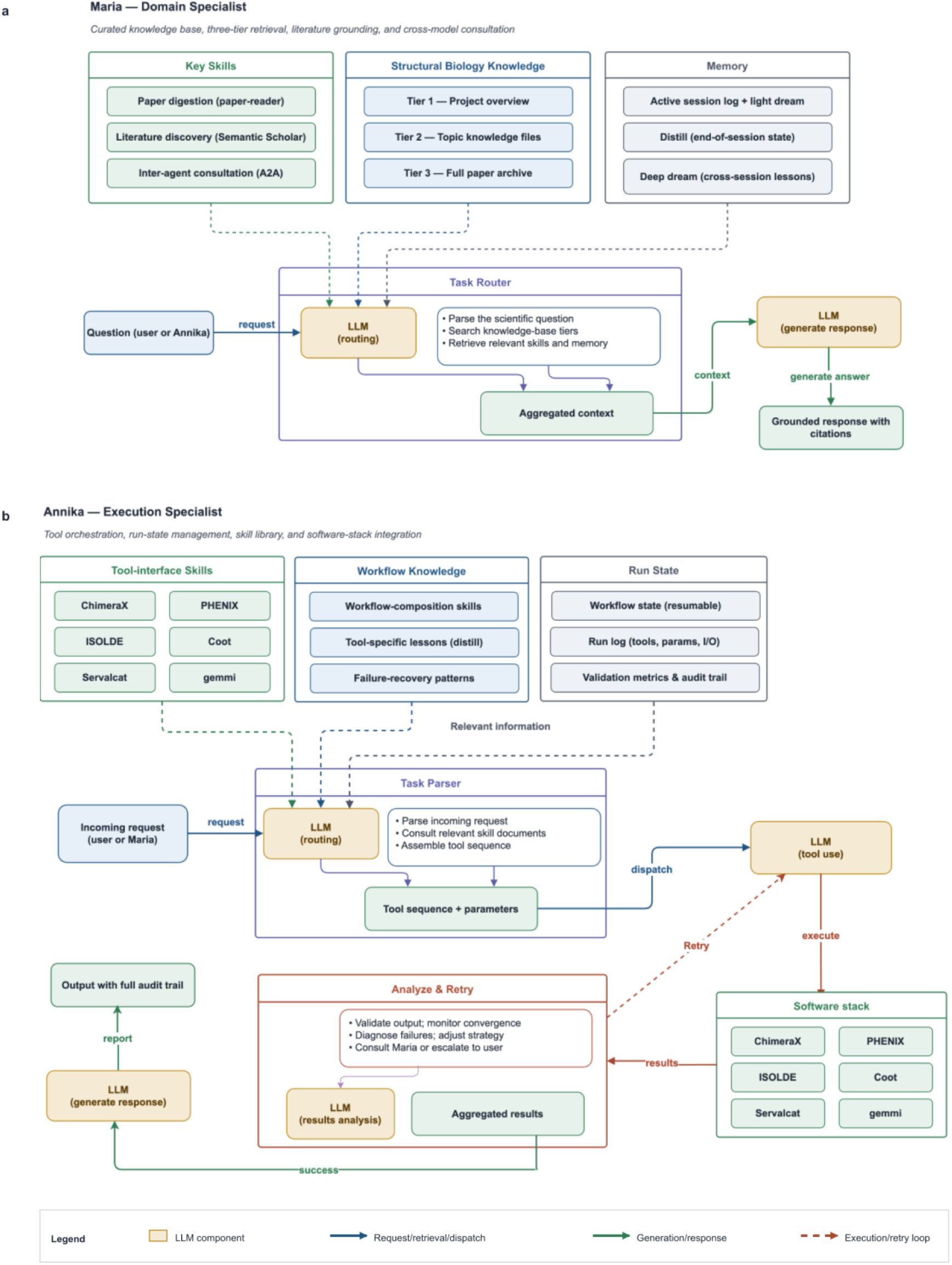
StructAgent agent architecture. **(a)** Maria’s domain-specialist architecture, including literature discovery, paper digestion, inter-agent consultation, the three-tier knowledge base and scientific memory. **(b)** Annika’s execution-specialist architecture, including tool-interface skills, workflow-composition skills, run state, validation metrics and retry logic. Approval gates are applied to strategy changes, model-changing corrections and deposition-related actions. Resource-release boundaries are described in the Data Availability and Code Availability sections.

**Fig. 6.**
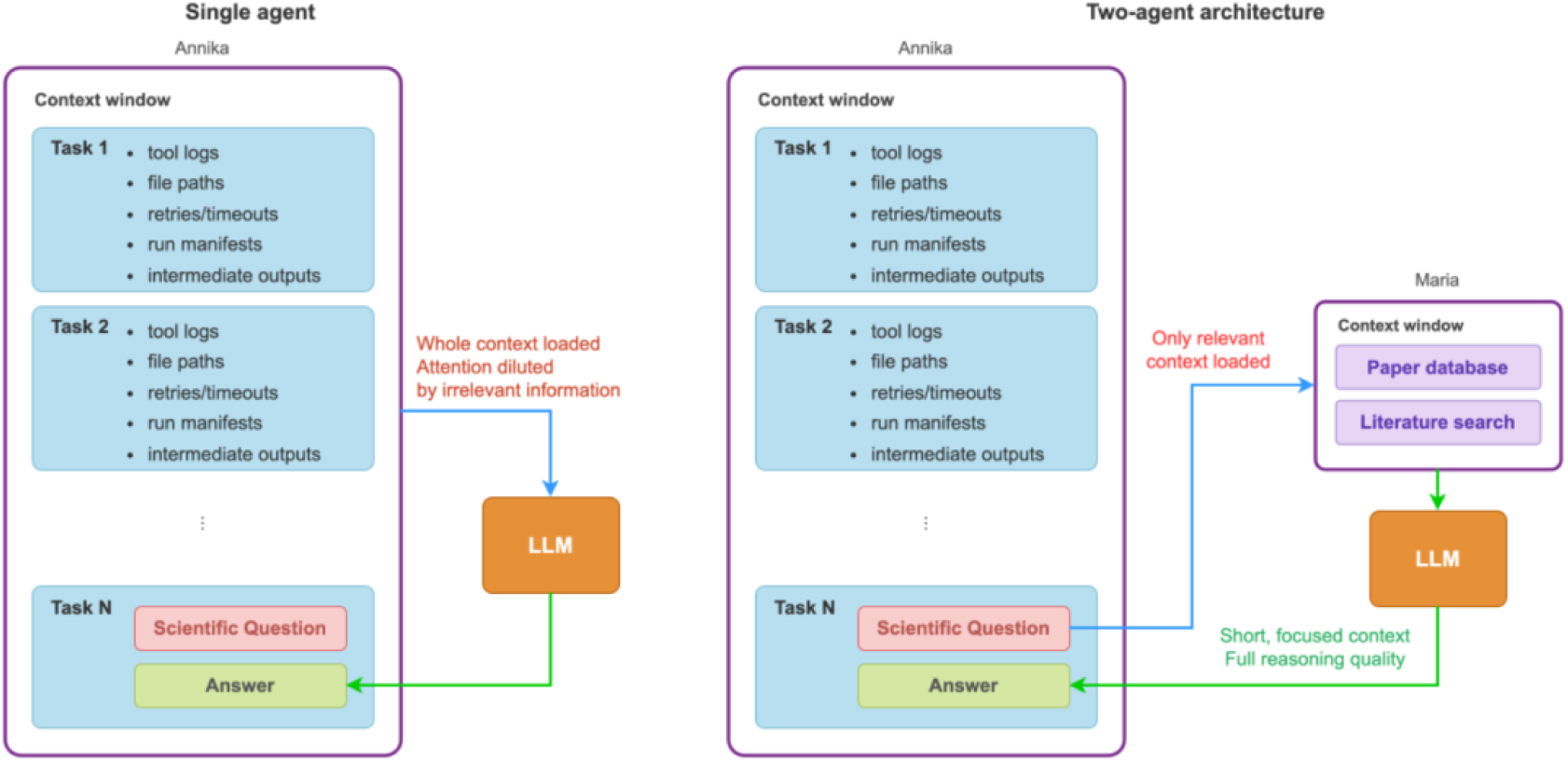
Context separation in single-agent and two-agent designs. In a single-agent design, accumulated execution detail competes with domain-level reasoning in the same context window. StructAgent keeps execution history in Annika’s session and sends focused scientific questions to Maria. This figure illustrates the design rationale; quantitative ablation of context separation was not performed.

## Discussion

StructAgent addresses a practical bottleneck in cryo-EM model building: not the absence of capable fitting, refinement or validation programs, but the difficulty of coordinating them across long, project-specific workflows. The system is therefore best understood as an auditable orchestration resource. It helps a structural biologist turn a modeling question into software actions, intermediate checks, recovery steps and evidence packages, while keeping model-changing decisions and final interpretation under expert control.

The three examples make complementary claims about this orchestration layer. The proteasome rebuild tests scale and expert handover: StructAgent coordinated a 64-chain rebuild from an earlier related template, improved all chains relative to the rigid-body template fit and produced a model suitable for targeted expert review. The ribosome example tests scalable strategy construction: Maria translated bond-valence and coordination chemistry into an audit specification, and Annika applied it to 530 metal sites, generated restraints and refined 75 user-approved corrections. The ongoing-work ligand example tests scientific reasoning: conventional tools could generate poses and density scores, but Maria connected residue chemistry, homologous folate binding and substrate-channeling logic to a defensible orientation choice, then defined a validation endpoint that preserved weak-tail uncertainty. Together, these examples show that the useful unit is not a single tool call, but a reviewable chain of reasoning, execution and validation.

StructAgent’s added value is the connection between established tools. ChimeraX, ISOLDE, PHENIX, Servalcat/REFMAC5, Coot, gemmi and validation servers perform the structural calculations. StructAgent links them into workflows, records how they were used, monitors intermediate outcomes, recognizes when downstream symptoms indicate broken assumptions and preserves repairs as reusable workflow knowledge. Across the case studies, all 14 tracked tool-level failures were recovered, but the count is less important than the form of the recovery: coordinate-frame shifts, force-field template gaps, restraint-format incompatibilities and refinement artifacts became inspectable events with causes, fixes and provenance, rather than hidden troubleshooting steps in a manual session.

This provenance addresses a broader documentation problem in structural biology. Published methods sections often name the software used for model building and refinement, but rarely capture the actual command sequence, parameter changes, failed branches, diagnostic checks or rationale that shaped the final model. StructAgent produces session records containing tool invocations, inputs, outputs, validation metrics, errors and recovery decisions, with software versions captured where available. These records do not make LLM-guided orchestration deterministic, and they do not remove the risk of a plausible but wrong recommendation. They make the workflow inspectable: a reviewer or future user can ask what was run, why a route changed, which alternatives failed and what evidence supported a model-changing decision.

Trust in StructAgent comes from constraint rather than autonomy. Atomic calculations are delegated to established structural-biology programs. Literature and skill documents constrain the reasoning space. Strategy changes, ligand-orientation choices, ion reassignments, residue mutations, broad refinement-route changes and deposition-related actions require user approval. These safeguards are not equivalent to formal verification, and they cannot guarantee that every recommendation is scientifically correct. They do, however, make the system closer to a disciplined and documented operator than to a black-box structure generator.

Skills are the reusable unit of this resource. A skill is not just a prompt fragment; it is a readable protocol that specifies expected inputs, command patterns, validation checks, common failure modes and recovery actions. Skills allow operational expertise to be revised after real use and reused in future sessions. The Maria-Annika separation is one implementation of this idea, designed to keep execution-heavy context, logs and file-state details separate from strategy, literature grounding and validation criteria. It is not a fixed requirement of the resource. The same skill documents could be used by a single sufficiently capable agent, by a different multi-agent arrangement, or by a general coding assistant that can read protocols and operate local files. Additional specialist agents, for example focused on biological interpretation, could be added without changing the principle that skills and logs are the portable substrate.

More generally, an agent does not know all experimental context: sample preparation, construct design, biochemical expectations, map-processing choices, unpublished controls and project priorities may be available only to the researcher. StructAgent can organize evidence and propose actions, but it should not be trusted blindly to decide what is biologically plausible or deposition-ready. The next step is therefore not to remove the expert from model building, but to reduce repetitive protocol execution. As more skills are generated, tested and revised, routine refinement setup, validation checks, simple fitting trials and repeated audit procedures should become increasingly agent-executable with limited intervention, while unusual chemistry, ambiguous density, weak local resolution and biologically consequential model changes continue to require explicit expert review. Future evaluation should combine public archives with installable code, sanitized skills, versioned command wrappers, smoke-test data and releasable case-study bundles; external installation by independent users; and prospective comparisons with manual or semi-automated workflows. StructAgent should not be presented as an autonomous deposition engine or a replacement for structural biology expertise. Its contribution is narrower and more practical: it preserves expert agency while making complex cryo-EM model-building workflows easier to execute, inspect, repeat and improve.

### Online Methods

### Overall system design

StructAgent is a domain-specific agent system for cryo-EM model building and refinement, built on a substantially modified version of the OpenClaw multi-agent framework (https://github.com/openclaw/openclaw). The base framework provides session management, tool execution and workspace state. StructAgent extends it with custom agent roles, memory handling, domain-specific inter-agent communication, structural-biology tool integration and session management for long, information-dense workflows.

The language model is embedded in an operational environment with explicit tools, bounded task protocols, persistent state and structured communication channels. The system is therefore used as a workflow controller rather than as a free-form chatbot. All structural-biology calculations are delegated to local software.

### Agent roles

Maria is the domain-specialist agent. She interprets scientific questions, evaluates strategies, grounds recommendations in literature and project memory, diagnoses domain-level failure modes and defines validation criteria. Her resources include a three-tier structural-biology knowledge base, literature-retrieval tools and project-specific memory.

Annika is the execution-specialist agent. She translates user intent or Maria-approved strategies into concrete software actions, sequences tool invocations, manages workflow state and handles error recovery. Annika’s tool layer covers ChimeraX, ISOLDE, PHENIX, Coot, Servalcat/REFMAC5, gemmi and supporting scripts. When a task spans multiple programs, Annika composes the relevant skills dynamically and records the run state.

The internal architectures of Maria and Annika are summarized in Fig. 5.

### Knowledge base

Maria’s curated knowledge base is organized in three tiers, designed for efficient retrieval within token-budget constraints.

**Tier 1 — project overview (approximately 500 tokens, always loaded).** A lightweight brief captures the current scientific focus, key open questions and recent findings. This is loaded at the start of every session, giving Maria immediate project context.

**Tier 2 — topic-specific knowledge files (approximately 1-3k tokens each, loaded on demand).** Distilled summaries are organized by topic: method comparison tables, resolution-dependent parameter guidelines, decision trees for common tasks and operational lessons. These encode synthesized, actionable rules, for example, “B-factor refinement should be deferred until geometry is stable” or “maps below 4 Å typically require tighter secondary-structure restraints”, rather than individual paper summaries. Each rule traces back to its source papers.

**Tier 3 — full paper archive (searched by script, never bulk-loaded).** Individual paper digests, structured metadata and a cross-reference index capture relationships between papers. The archive currently contains more than 80 papers and is queried through targeted searches. A full inventory is provided in Supplementary Table 2.

This three-tier design keeps Maria’s token cost approximately flat as the knowledge base grows: adding more papers enriches Tier 3 and occasionally distills new rules into Tier 2, while the always-loaded context remains small.

When a question falls outside the curated database, Maria can extend retrieval through a literature-discovery skill that queries the Semantic Scholar API [33], deduplicates results against the existing database and triages candidates for relevance. New papers can be digested during a session or queued for later reading. The goal is to ground recommendations in curated or retrievable evidence rather than unsupported model memory.

### Skills

Skills are human-readable protocol documents that define how a class of tasks should be performed: required inputs, tool commands, parameter choices, validation checks, expected outputs and common failure-recovery procedures. Because skills are readable and versioned, they convert tacit operational expertise into a reusable and auditable resource.

### Skill architecture

StructAgent uses a dedicated skill-creation pipeline with a top-level classifier that identifies the type of skill needed and applies the appropriate architecture. **Tool-interface skills** (for example, ChimeraX, PHENIX and ISOLDE) encode software-specific knowledge: command syntax, parameter conventions, input/output formats and known quirks. **Workflow-composition skills** (for example, structural-build and structural-strategy) encode higher-level decision logic: which tools to use for which problems, in what order and with what validation gates. **Knowledge-acquisition skills** (for example, paper-reader and discovery) encode processes for building and maintaining the knowledge base itself.

### How skills are built

Tool-interface skills were assembled from official documentation, methodology papers, tutorials, observed command behavior and task logs, then revised after use on real projects. The resulting skill document is tested against real tasks, and failures drive iterative revision. Skills evolve through reviewed operational experience rather than through autonomous self-modification.

### Skill evolution through use

The ISOLDE skill provides a concrete example. The initial version could launch ISOLDE sessions but had no robust way to monitor progress, detect convergence or catch failures; GUI state changes were invisible to the agent. Through iterative debugging across real tasks, the skill acquired (i) a Python monitoring bridge that reads simulation state from a running ISOLDE session at fixed intervals; (ii) explicit input-file verification, checking atom naming, chain assignments and residue completeness before launch, after discovering that ISOLDE fails silently on certain input errors; (iii) a pre-launch model-preparation checklist; and (iv) convergence criteria calibrated to different resolution ranges. At the end of sessions, the system can summarize completed work and propose reusable lessons; deeper lesson-distillation steps extract candidate updates to skills or knowledge files, such as new preflight checks, failure signatures or parameter rules. These updates are stored as proposed changes and presented for user review before being applied; the system does not autonomously modify its own protocols.

A full skill inventory for both agents is provided in Supplementary Table 1.

### Execution tracking and inter-agent communication

During multi-step tasks, Annika records tool invocations, software versions where available, input files, command-line parameters, standard output and error streams, output files and validation metrics, including MolProbity-based geometry checks [34]. These logs allow interrupted workflows to be resumed, prevent repeated failed approaches and provide provenance records for completed runs.

A worked example is ISOLDE convergence monitoring. ISOLDE is an interactive molecular-dynamics flexible-fitting tool designed for human use in a 3D graphical environment. Making it agent-controllable required bridging the gap between the LLM’s text-based reasoning and a GUI tool that assumes visual feedback. StructAgent uses Python monitoring scripts that read simulation state from the running ISOLDE session at fixed intervals, extract quantitative metrics (CC_mask, geometry statistics, simulation energy), detect stalls or failures and report structured status back to the agent. The agent can therefore launch an ISOLDE session, monitor its progress through numbers rather than visual inspection and terminate refinement when the change between consecutive CC_mask readings falls below a convergence threshold. The same pattern, a polling bridge that converts GUI state into structured status, is reused by other interactive tools in Annika’s stack.

Long model-building sessions can accumulate enough operational detail to interfere with scientific reasoning. StructAgent addresses this with context compaction adapted from Lossless Context Management [35], by keeping execution history in Annika’s context and by sending focused scientific questions to Maria only when domain reasoning is needed (Fig. 6). Inter-agent communication is implemented through the Agent-to-Agent (A2A) protocol [36], adapted with sender-side logging, long-running consultation timeouts and structured messages that separate scientific questions from operational metadata. Because Maria and Annika can run on different LLM backends, and Maria in particular can draw on more than one model, inter-agent consultation also functions as a form of cross-model reasoning, bringing complementary strengths to bear on the same scientific question.

### Memory architecture

StructAgent uses a multi-timescale memory architecture: active session memory, periodic context compaction, end-of-session reflection and lesson distillation.

### Active session memory

During a session, every action, tool output, decision and inter-agent consultation is recorded in a structured session log. This provides the operational history for the current task, supports backtracking and forms the raw material for downstream memory processes.

### Periodic context compaction

Long cryo-EM modeling sessions can run for hours, accumulating more context than the LLM’s active processing limit can usefully hold. StructAgent uses periodic context compaction, an LLM-based process that summarizes completed portions of the session while preserving key decisions, metrics and outcomes; the original, uncompacted history remains stored and can be retrieved when needed. This mechanism builds on the Lossless Context Management (LCM) system [35], with custom compaction strategies tuned for the information density of structural-biology sessions, where a single tool output may contain hundreds of lines of validation statistics, only a few of which matter for future decisions.

### End-of-session reflection (session distill)

At the end of each working session, the system runs a structured reflection step: it reviews what was accomplished, what problems were encountered, what decisions were made and what remains to be done, and writes a compact working-state file that the next session loads on startup. This is structured synthesis rather than automatic compaction, closer to a researcher writing up lab notes at the end of the day.

### Lesson distill (cross-session knowledge distillation)

Periodically, triggered by the user after significant sessions, the system performs a deeper reflection that extracts reusable lessons from accumulated operational experience and proposes updates to skills, knowledge-base entries or decision rules. These proposals are presented to the user for review before being applied; the system never modifies its own skills or knowledge autonomously. Together, session distill and lesson distill form a closed loop: operational experience generates lessons, lessons improve skills and improved skills produce better operational outcomes.

### Per-agent memory specialization

Maria and Annika use the same memory infrastructure but store different kinds of information. Maria’s memory is primarily scientific: the three-tier knowledge base, evaluated literature, project-level decision context and accumulated domain rules. Annika’s memory is primarily operational: run history, workflow state, environment-specific behavior (which software versions are installed, which parameter combinations have been tested, which file formats each tool expects) and tool-specific lessons learned. This separation reflects the agents’ different roles and ensures that each agent’s persistent memory is optimized for its function.

### Language model backends

In the current deployment, both agents use Anthropic Claude Opus 4.6 [37] as the primary model for multi-step tool use and long-context reasoning. Maria can additionally use OpenAI GPT-5.4 [38] for complementary scientific consultation. Lightweight operational tasks, including session summarization, context compaction and structured logging, use Qwen 2.5 7B [39] locally. Backend assignment is a configuration parameter rather than an architectural requirement, and model identifiers and settings are recorded in session logs where available.

### Deployment and approval gates

The current deployment runs on a single Apple Mac mini (M4, 16 GB unified memory). Agent sessions, tool execution, inter-agent messaging and structural-biology software run locally on macOS with Apple Silicon. Primary LLM inference uses external API providers; the local machine does not require a GPU cluster. The user interacts through the OpenClaw messaging layer, including Telegram, WhatsApp or other chat clients.

The deployed structural-biology stack included ChimeraX [40], ISOLDE [7], PHENIX [5], Coot [41], Servalcat [8]/REFMAC5 [9], gemmi and task-specific scripts.

Approval gates are applied to strategy-level changes, ligand-orientation choices, ion reassignments, residue mutations, broad refinement-route changes and any deposition- or release-related action. Annika may run preflight checks, format conversions, validation scripts and retry attempts that do not change scientific model content. Major model-changing actions are presented to the user for review before execution.

Agents generated plans, commands, diagnostic summaries and candidate evidence packages, but did not autonomously decide final biological interpretations or deposition-ready coordinates. The author reviewed manuscript claims, case-study outputs, figure content and model-changing decisions. Tool permissions were limited by the local operating environment and by explicit approval gates; potentially destructive file operations, model-changing edits and release actions required human approval.

### Case-study validation protocols

For Example 1, validation used Phenix 2.0 (phenix.validation_cryoem, phenix.molprobity, phenix.mtriage and phenix.emringer) at 3.8 Å against the deposited EMD-8332 full map; half-maps were not used. PDB 5T0C was excluded from pipeline stages and used only for post hoc comparison. AlphaFold DB v4 monomers [26] were used for all 32 unique subunits.

ProSMART [25] distance restraints were generated from AlphaFold-predicted structures for Servalcat refinement, and AlphaFold monomers were used as reference models in PHENIX real-space refinement.

For Example 3, ligand restraints for 5-formimidoyltetrahydrofolate were generated with PHENIX/eLBOW. The Site 2 transferase active-site pose was transferred from PDB 1QD1/FON after structural superposition. Site 1 at the inter-domain interface was evaluated by residue-environment mapping, dual-orientation ChimeraX fitmap scoring and per-group density analysis on unrefined seed poses. Flexible fitting used ISOLDE; after the initial OpenMM template failure, a USER_LIG force-field XML was generated through antechamber, tleap and ParmEd. Final refinement used PHENIX real-space refinement with individual isotropic ADP and custom pterin planarity restraints. Validation included phenix.mtriage, per-group ligand density, B-factor parity and HOLE v2.3.2 channel profiling. The map and model files are withheld because the biological structure is from an ongoing study.

For Example 2, Annika extracted coordination shells for all 530 deposited metal positions in PDB 8B0X with gemmi, computed BVS values for user-specified candidate ions and assigned confidence grades from BVS, coordination number and geometry. Corrected sites were refined with REFMAC5 through Servalcat using bond-distance restraints and angle-derived 1–3 distance restraints. CMM validation was run on the original 530 positions; after correction, the 9 sites reclassified as water were excluded from corrected ion-score totals, leaving 521 ion sites.

### Data Availability

The public structural data used in this study are available from public repositories: EMD-8332 (EMDB), PDB 5GJR, PDB 5T0C, PDB 1QD1 and PDB 8B0X (Protein Data Bank). The cryo-EM map, model and accession information for the ongoing-work ligand example are from an ongoing study and are withheld from the initial public resource archive; they can be included in a confidential reviewer-only evidence bundle if requested and will be deposited separately upon publication of that work. Derived models, workflow traces and case-study evidence for Examples 1-3 are organized as reviewer-only evidence bundles for validation of the manuscript claims, not as public tutorial datasets in the initial public archive. Because the case studies demonstrate StructAgent workflows rather than report newly determined biological structures, formal PDB deposition of derived outputs was not pursued.

### Code Availability

The StructAgent source repository is available at https://github.com/bhgtiger/StructAgent under an Apache-2.0 license. The manuscript corresponds to commit cdbf720dbb19061aa94c10e7e52e329b54982854. A DOI-bearing Zenodo archive has been reserved for the submitted release: 10.5281/zenodo.20015169; reviewer access to the draft archive is available at https://zenodo.org/records/20015169?preview=1&token=eyJhbGciOiJIUzUxMiJ9.eyJpZCI6Ijc3 ZjkzNWNmLWQzY2EtNDI3Ny1iYjg4LTQ0NDEzZDBkZjg0YyIsImRhdGEiOnt9LCJyYW5k b20iOiI0ODYyNmM2MmU0Zjg4MTQ4N2RlZDAwN2M5OWY4NjY0MyJ9.mj-JUwE-vcdgzofuSQHZdUy_h7KhitwiZxBj0IEBXBLII8qfSXqJhK9Ns2ltbErxwnPAO2a5SJsZJU66du7 J7Q.

## Acknowledgements

We thank colleagues in the B8 group at the Netherlands Cancer Institute for constructive feedback during development. We are grateful to Henri van Luenen for supporting and encouraging AI-assisted research within the group. We also thank Xuli Wang for her support throughout this project.

## Author Contributions

X.G. conceived the project, designed the system architecture, implemented StructAgent, conducted all experiments, analyzed the data and wrote the manuscript. Large language model assistants were used for manuscript drafting and editing under the author’s direction and review.

## Supplementary Tables

**Supplementary Table 1.**
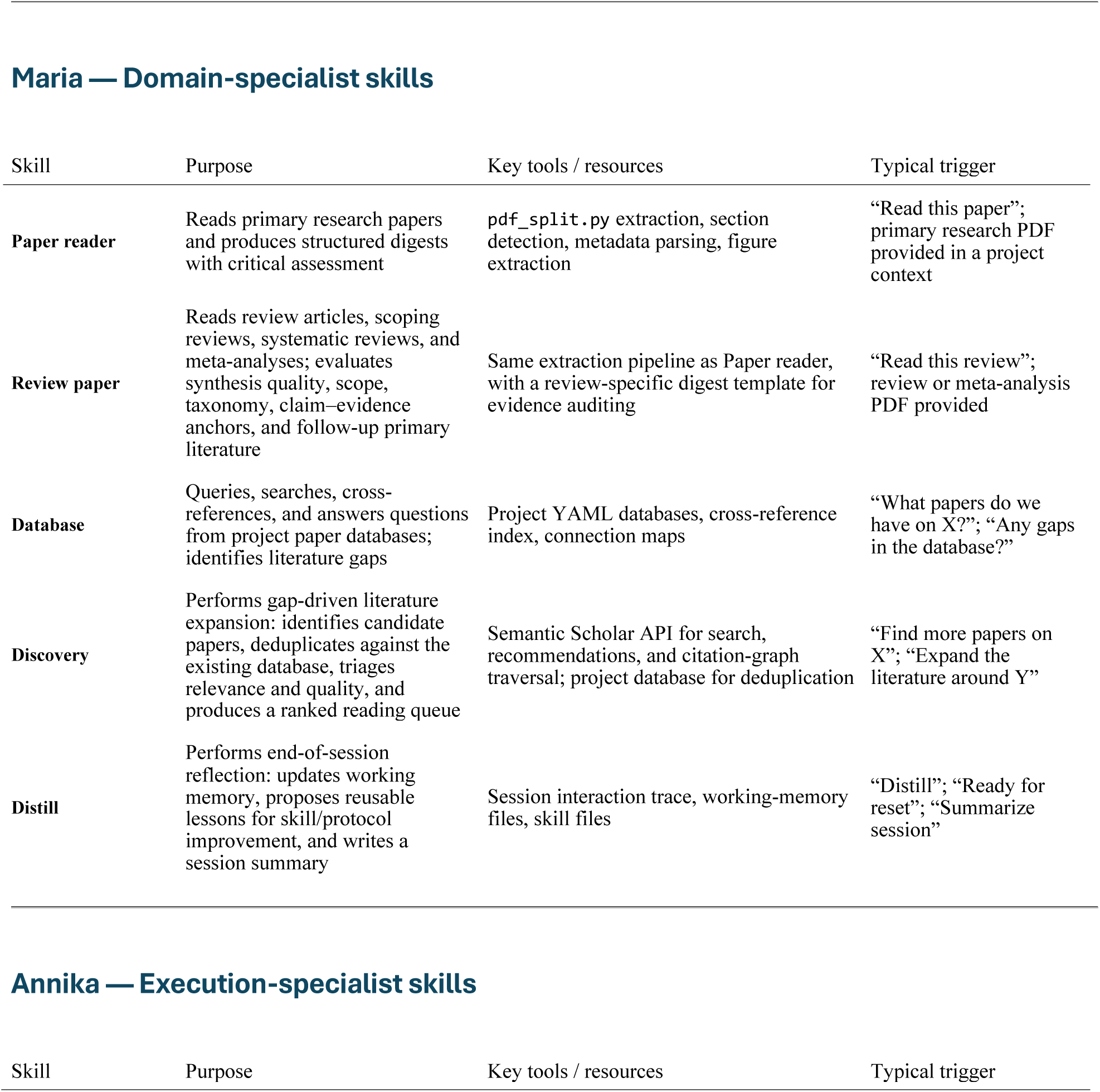
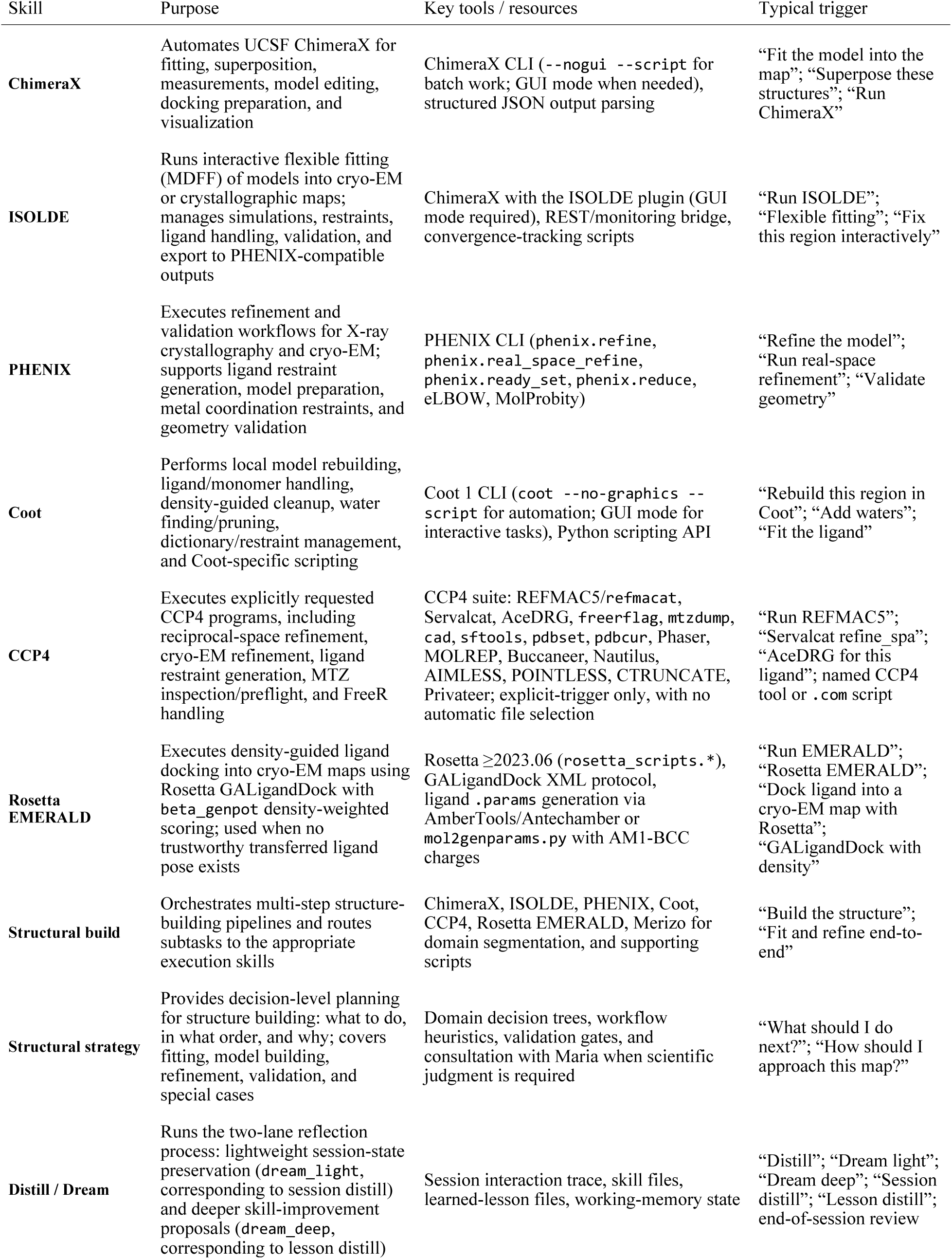
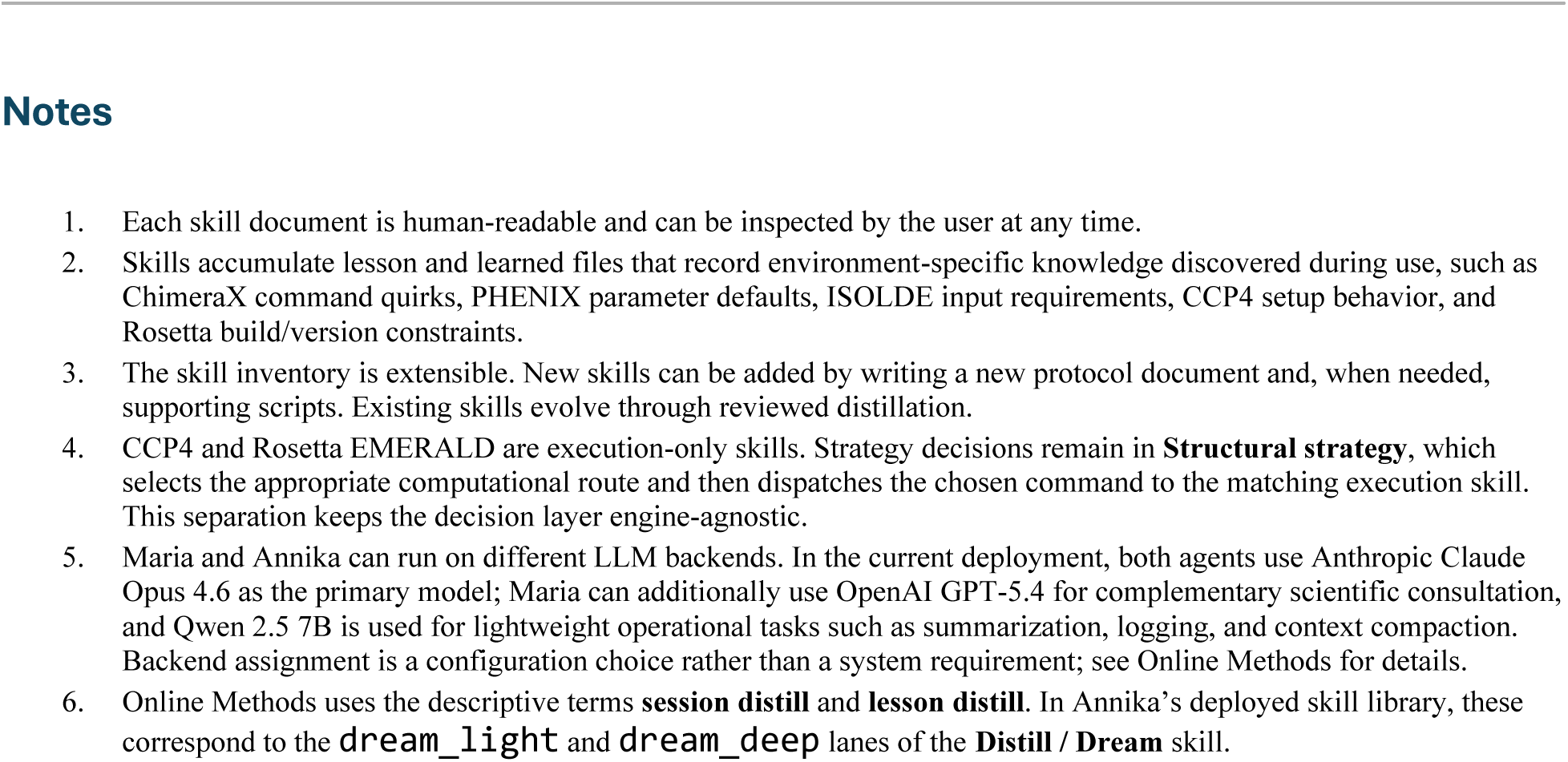
StructAgent skill inventory. Skills are structured protocol documents that define how each agent handles a class of tasks. Each skill specifies the task scope, required tools, execution steps, validation criteria, and failure-recovery logic. Skills evolve through the reviewed distillation mechanisms described in Online Methods.

**Supplementary Table 2.**
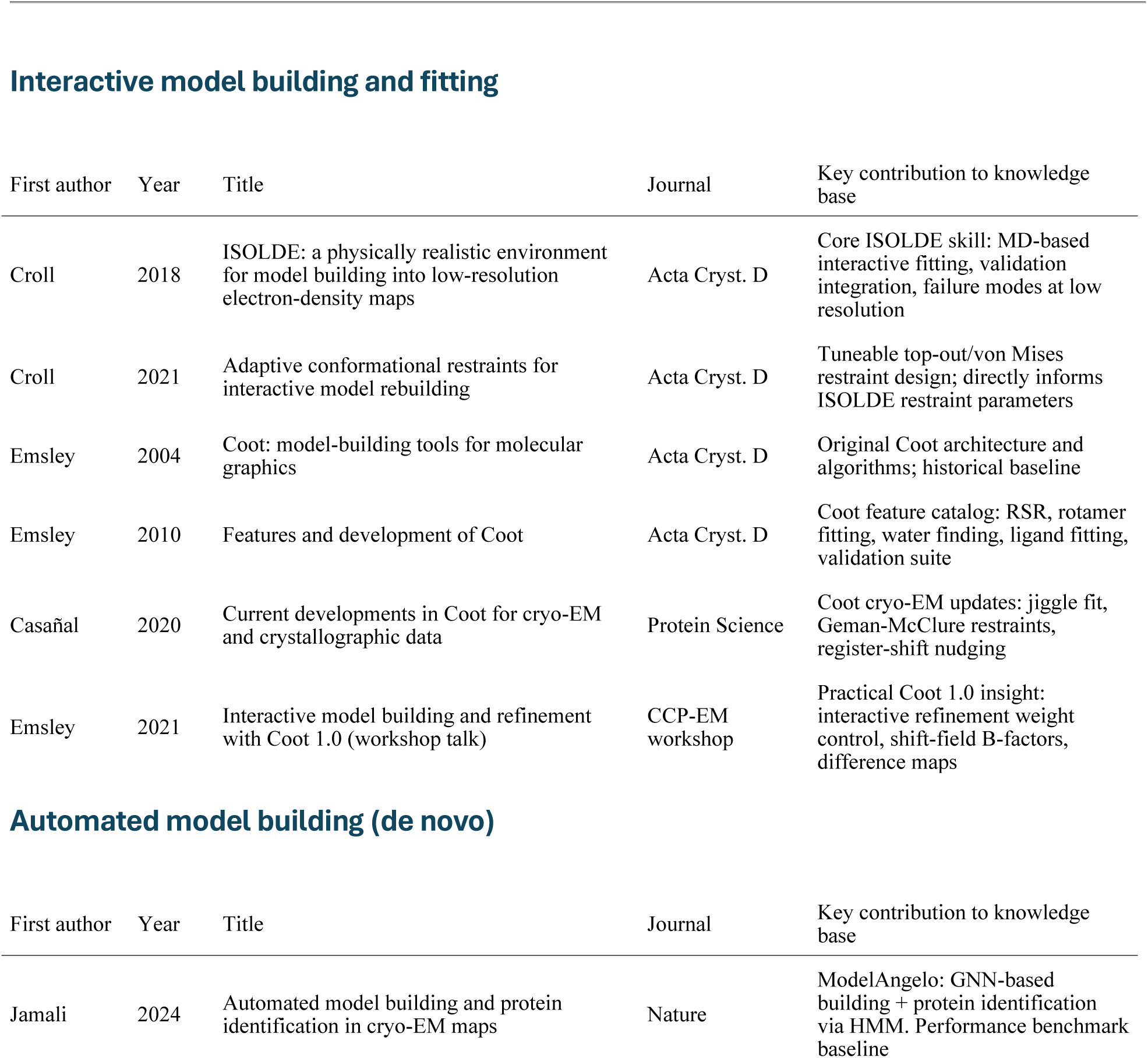
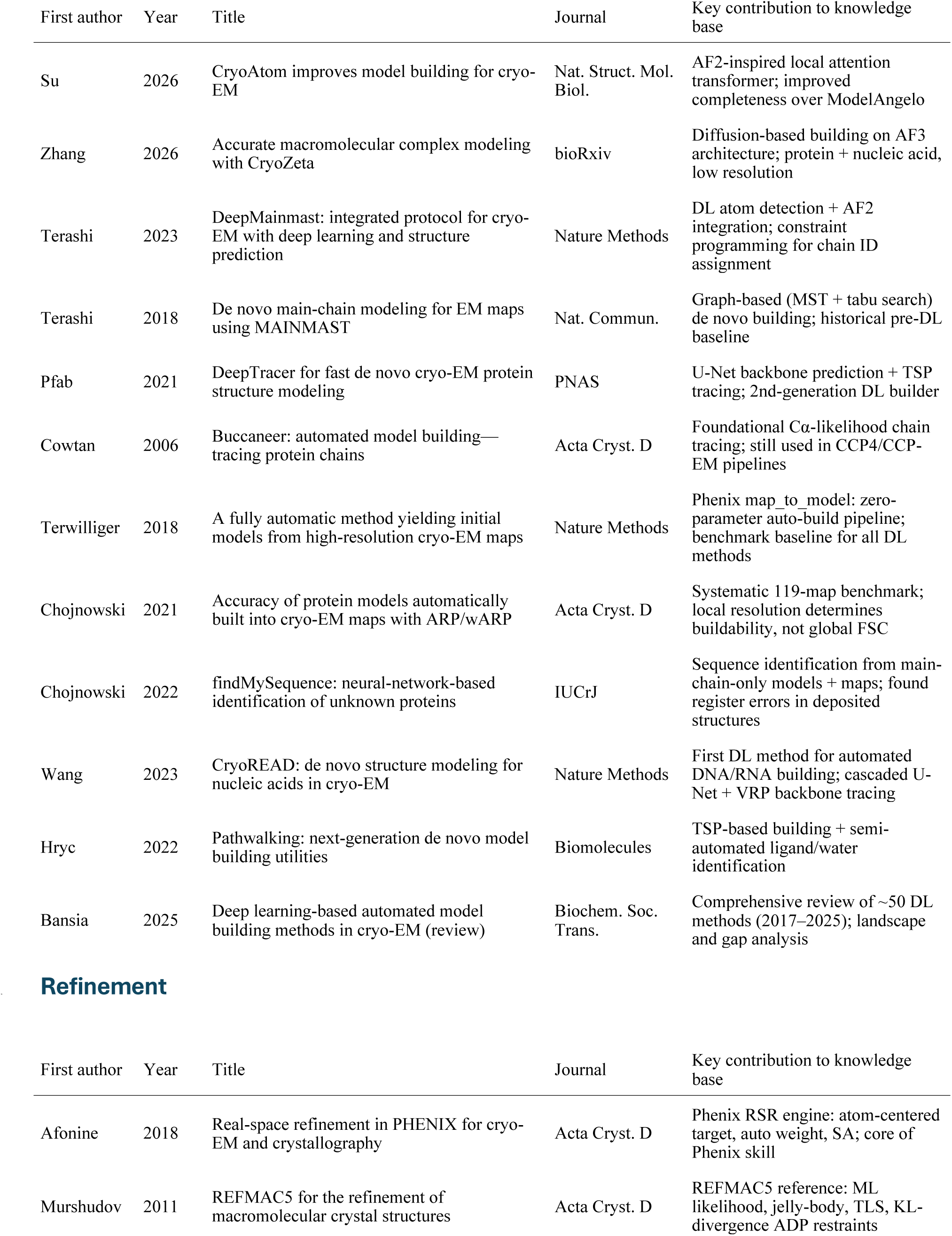
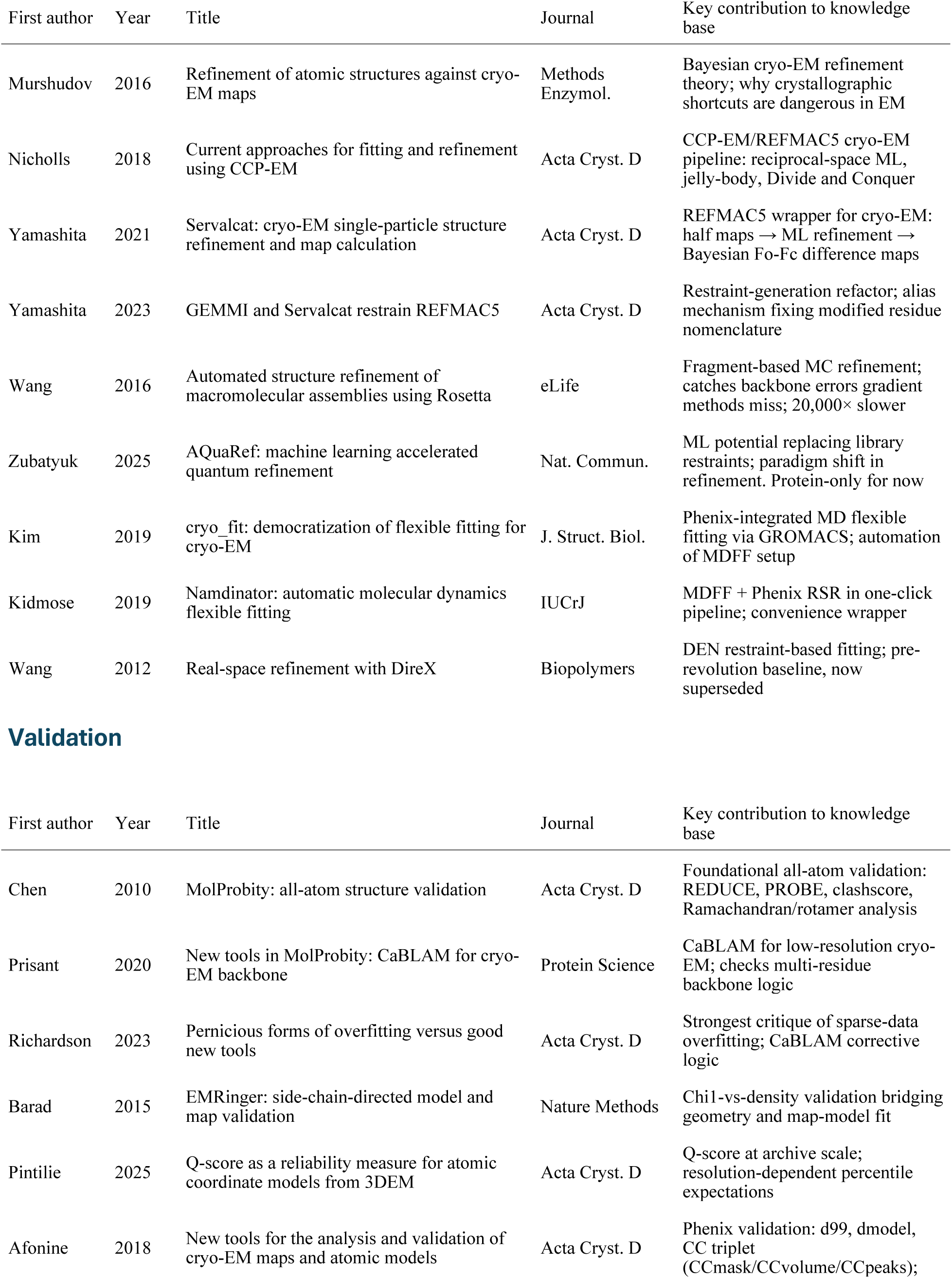
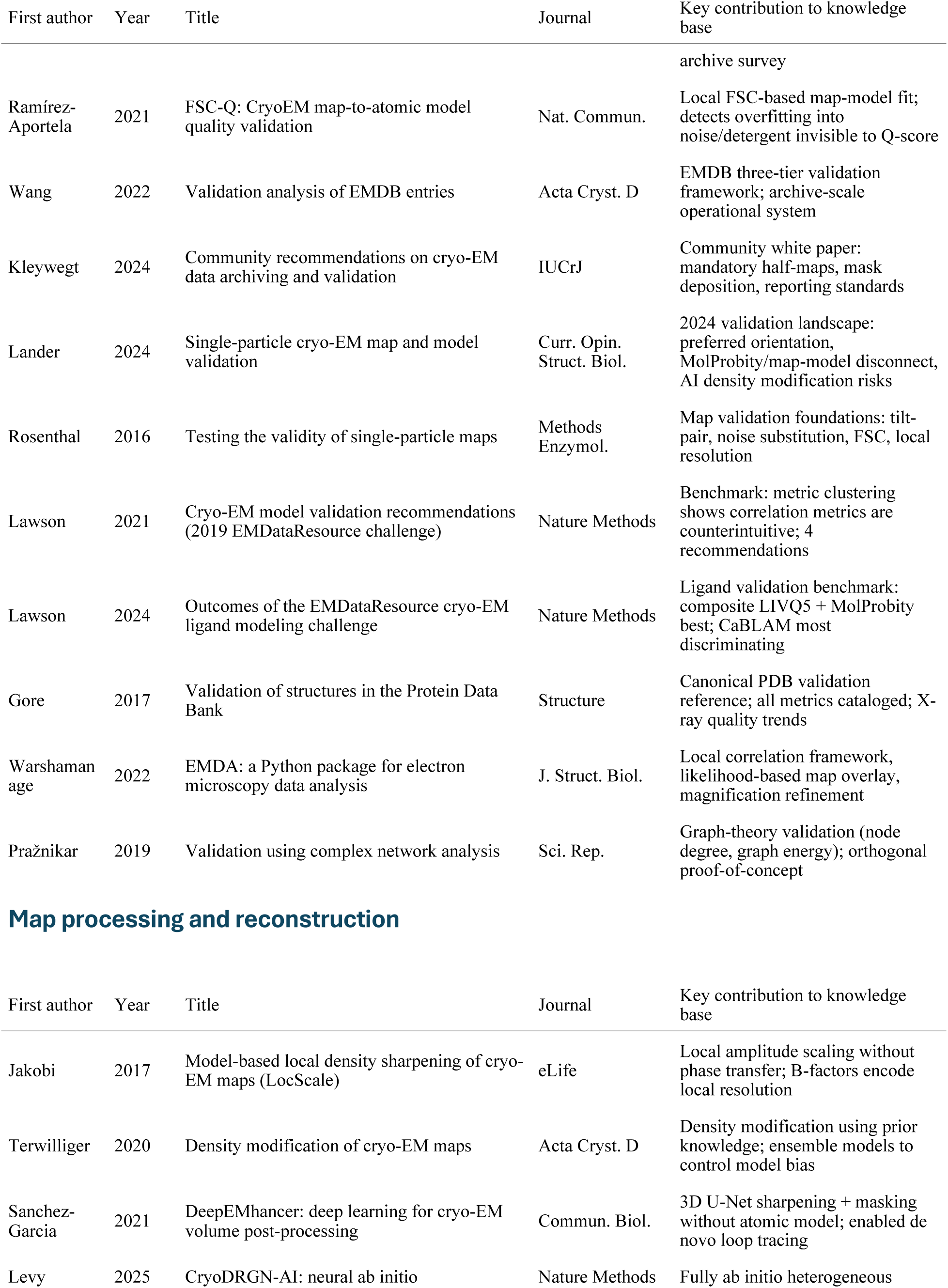
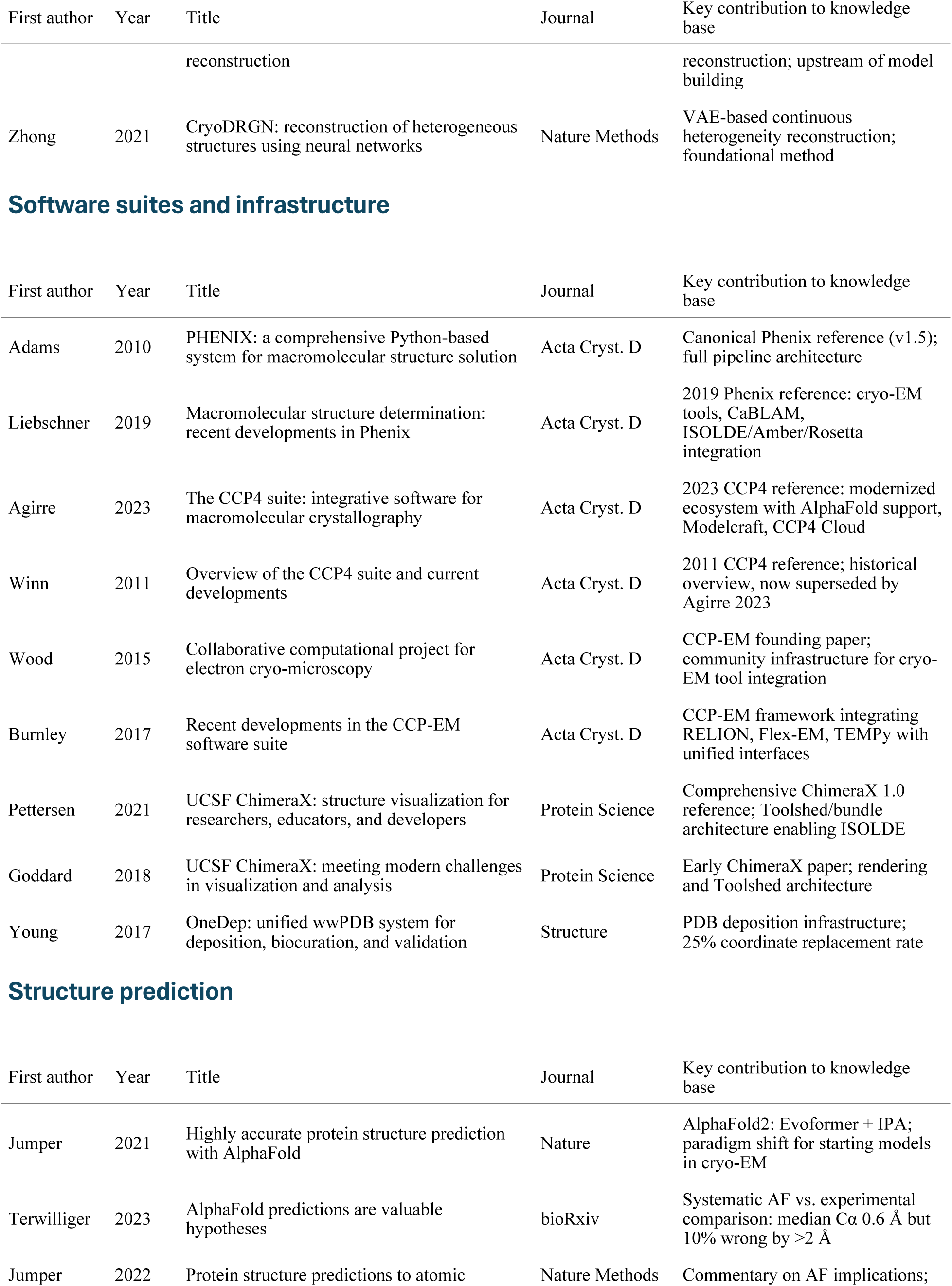
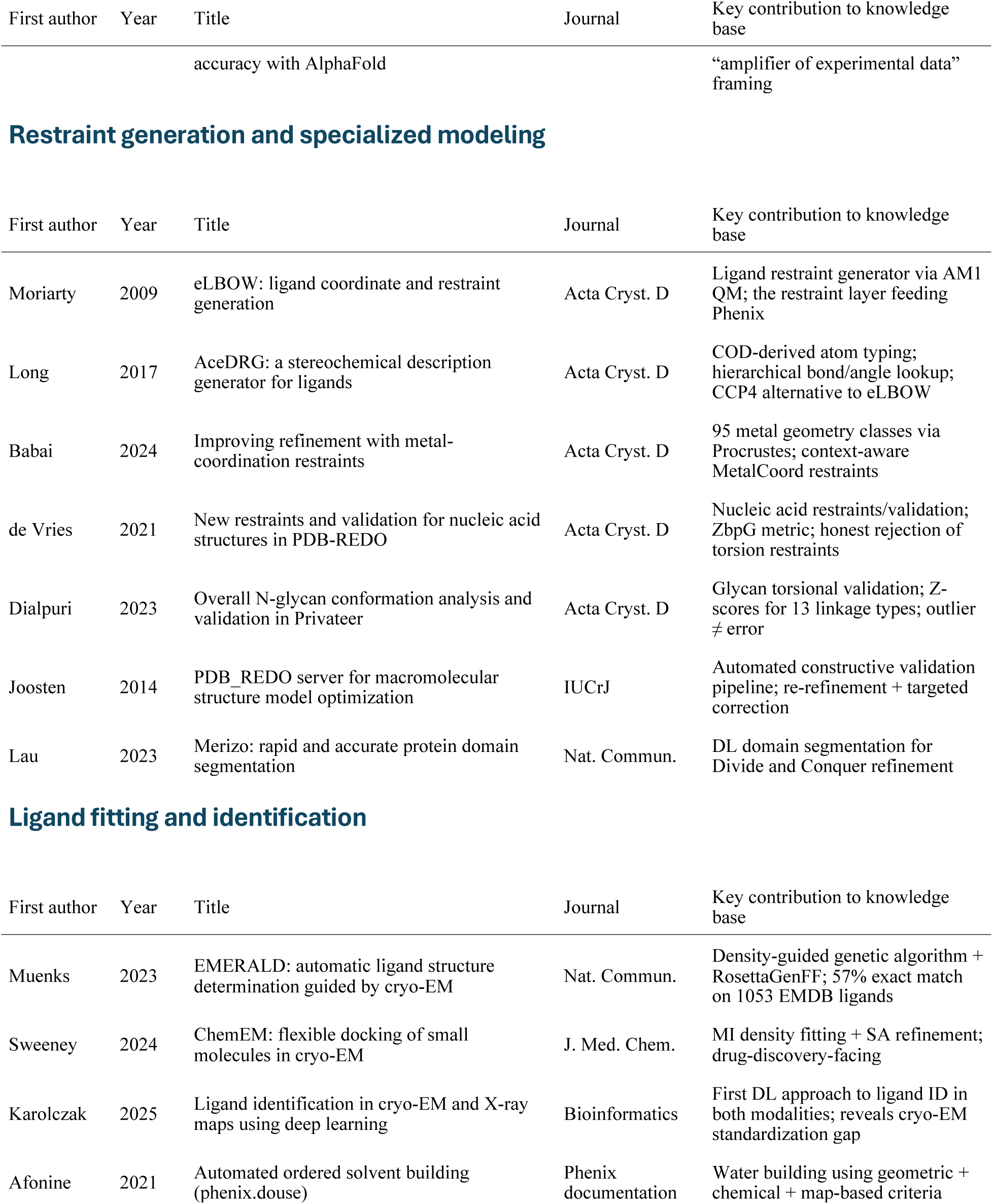
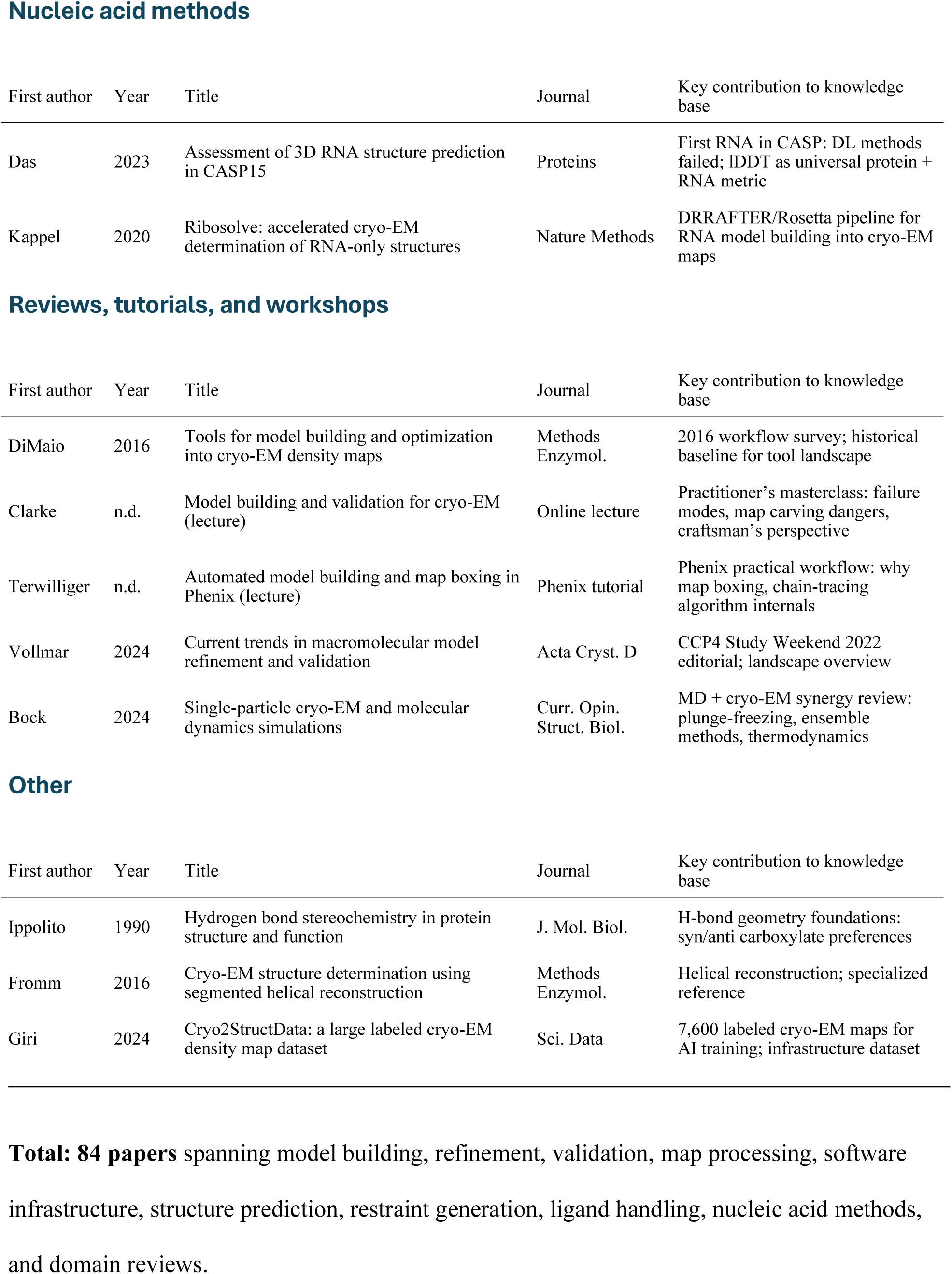
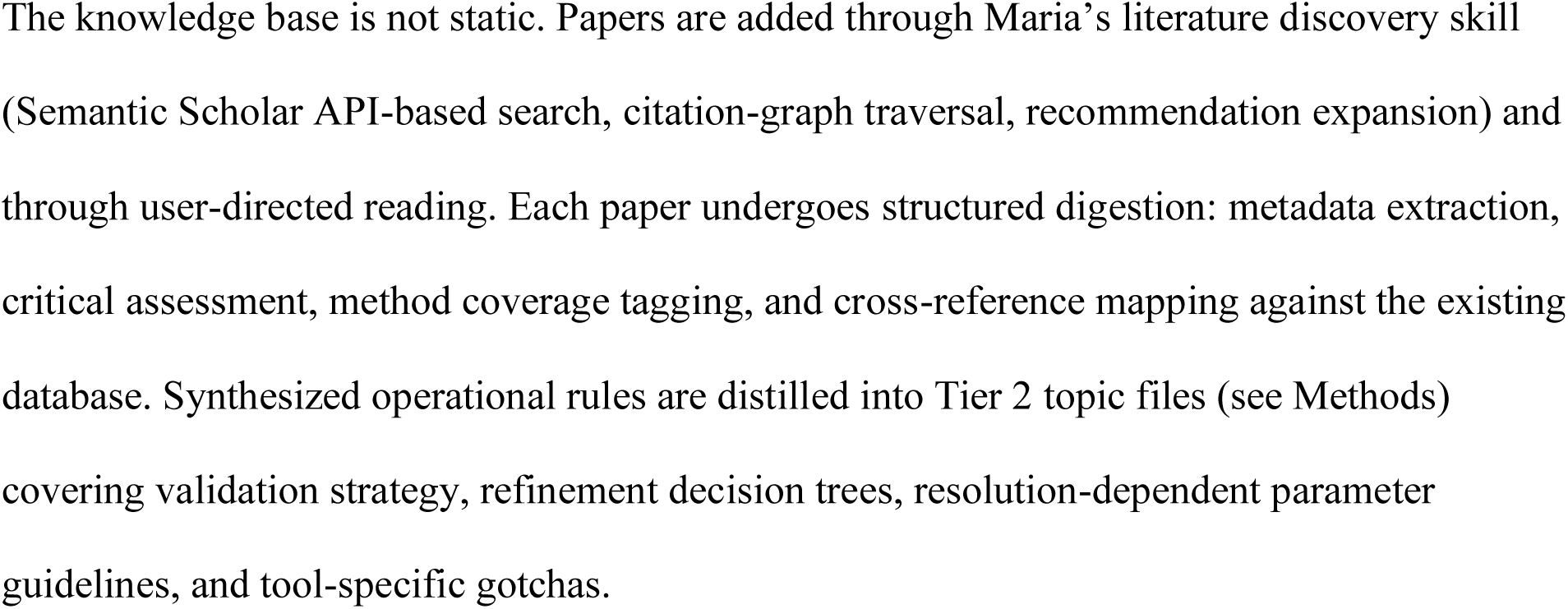
Knowledge base paper inventory. This table lists all papers currently in StructAgent’s curated knowledge base, organized by category. Papers are stored as structured YAML entries with metadata, structured digests with critical assessments, and cross-reference connections. The knowledge base feeds Maria’s three-tier retrieval system (see Methods): Tier 1 (project briefs) and Tier 2 (synthesized topic files) are distilled from these sources; Tier 3 provides searchable access to individual digests and metadata.

**Supplementary Table 3.**
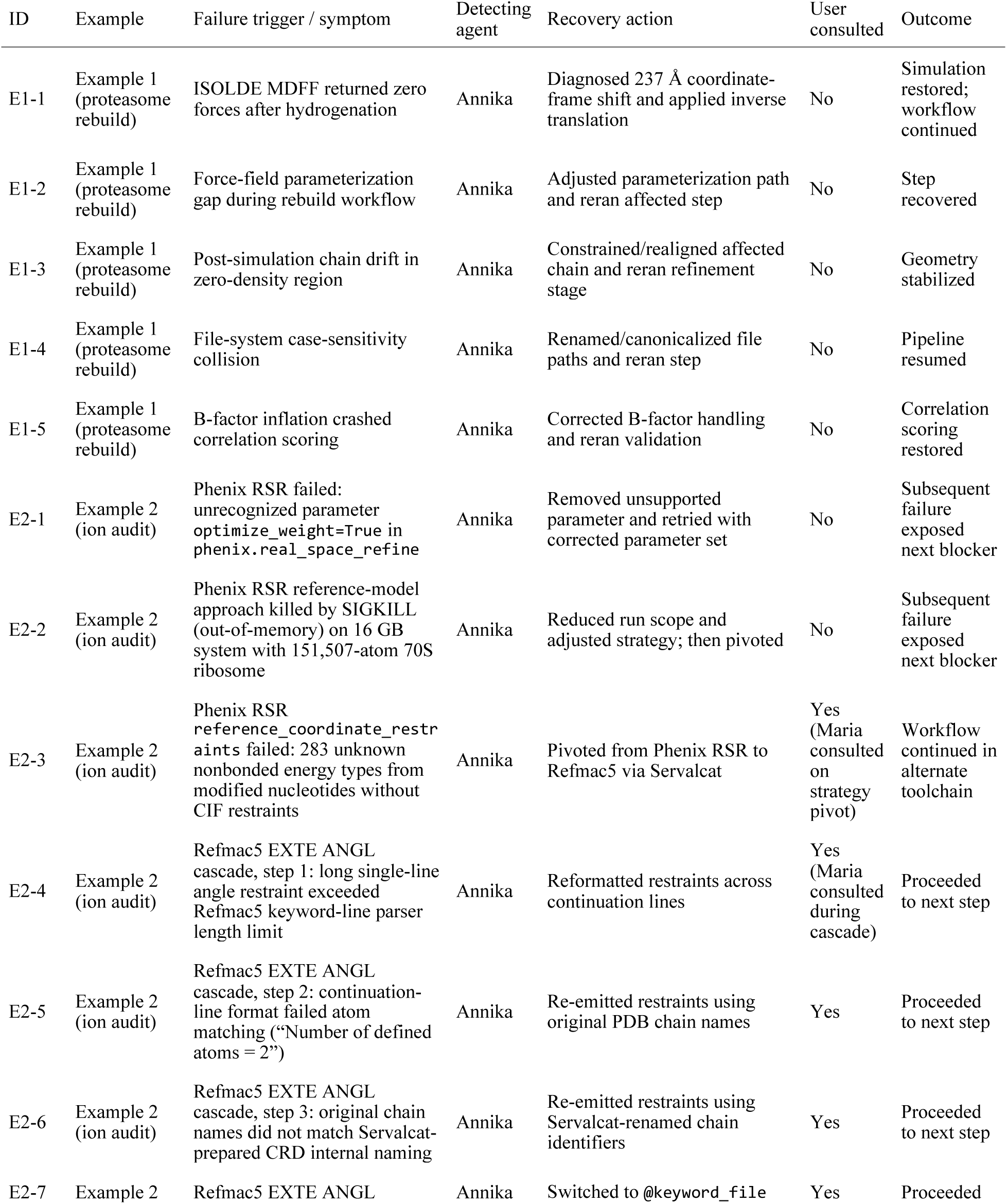
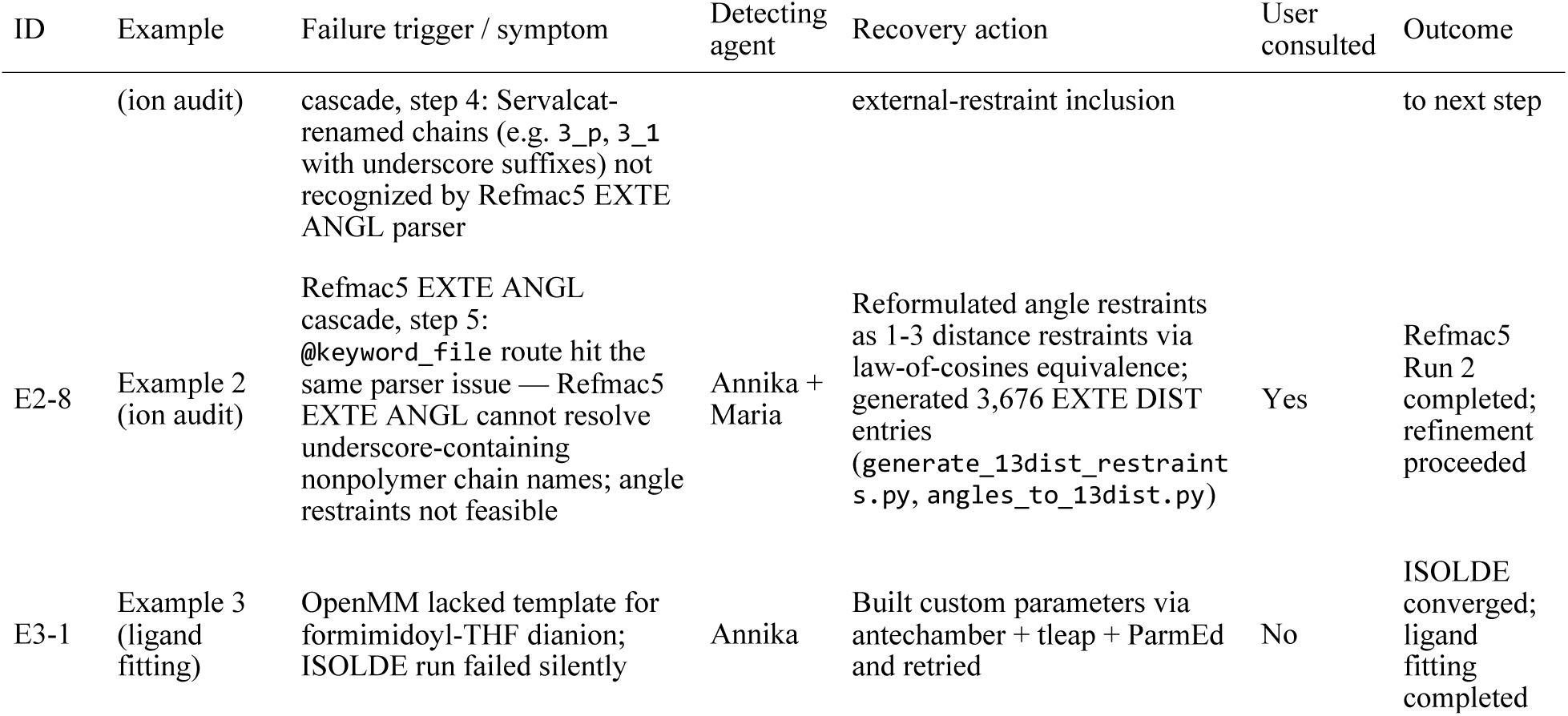
Tool-level failure and recovery events across the three case studies.

